# Near atomic structure of the inner ring of the *Saccharomyces cerevisiae* nuclear pore complex

**DOI:** 10.1101/2021.11.26.470097

**Authors:** Zongqiang Li, Shuaijiabin Chen, Liang Zhao, Guoqiang Huang, Xiong Pi, Shan Sun, Peiyi Wang, Sen-Fang Sui

## Abstract

Nuclear pore complexes (NPCs) mediate bidirectional nucleocytoplasmic transport of substances in eukaryotic cells. However, the accurate molecular arrangement of NPCs remains enigmatic owing to their huge size and highly dynamic nature. Here we determined the structure of the asymmetric unit of the inner ring (IR monomer) at 3.73 Å resolution by single-particle cryo-electron microscopy, and created an atomic model of the intact IR consisting of 192 copies from 8 subunits. In each IR monomer, two approximately parallel rhomboidal structures of the inner and outer layers are sandwiched with the Z-shaped Nup188-Nup192 middle layer and Nup188, Nup192 and Nic96 link all subunits to constitute a relatively stable IR monomer, while the intact IR is assembled by loose and instable interactions between IR monomer. These structures reveal various interaction modes and extensive flexible connections in the assembly, providing a structural basis for the stability and malleability of IR.

**One-Sentence Summary:** Accurate structure of inner ring reveals its assembly and function in dilation and constriction of nuclear pore complex.

## Main Text

The presence of nucleus, as the most distinguished hallmark between eukaryotes and prokaryotes, divides a eukaryotic cell into isolated compartments. Macromolecular transport in and out of the two-membraned nuclear envelope (NE) is essential for spatially separated transcription and translation ^1–3^. Nuclear pore complexes (NPCs), embedded in NE, are the massive bidirectional transport channels between the nucleus and the cytoplasm ^4^. Since the first glimpse about seventy years ago ^5,6^, NPCs have attracted increasing attentions from researchers. NPCs are the largest proteinaceous assembly in vivo, which consist of more than 550-1000 subunits from about 30 different nucleoporins (Nups) with the molecular mass of 60-120 megadalton from yeast to human, and are the sole gatekeepers controlling the nucleocytoplasmic transport of macromolecules ^7–9^. Perturbations of NPCs are known to cause numerous diseases, including viral infections, developmental diseases, neurodegenerative diseases and cancers ^10–14^.

It has always been a challenging task to comprehensively understand the elaborate structure and formidable function of NPC giving its huge size, complexity, flexibility and highly dynamic nature ^15,16^. In recent years, profited from the innovation of cryo-electron microscopy (cryo-EM) technology and the development of protein purification method, several teams have acquired the intact NPC samples successfully and determined the three-dimensional (3D) maps at 20-30Å resolution, including *Saccharomyces cerevisiae* NPC (*sc*NPC) ^17,18^, *Chlamydomonas reinhardtii* NPC (*cr*NPC) ^19^, *Xenopus laevis* NPC (*xl*NPC) ^20–22^ and *Homo sapiens* NPC (*hs*NPC) ^23,24^. However, owing to the barrier of low resolutions, the accurate molecular arrangement of NPC subunits remains thus far enigmatic.

Earlier analyses of protein-protein interactions and mass spectroscopy (MS) studies suggested that NPC can be divided into several stable subcomplexes, including the well-established Y complex ^25–27^, the scaffold inner ring complex ^24,28^, the CNT complex (channel nup trimer) ^28,29^, cytoplasmic 82 complex ^30,31^, nucleoplasmic basket complex ^32^ and the transmembrane ring complex ^17,33^, which further oligomerize into a cylindrical structure. Scientists have been devoted continuously to solve these structures of subcomplexes or subunits in an attempt to achieve the complete NPC structure as a jigsaw puzzle ^24,25,28,29,31^. However, so far only some fragments of subunits and partial structure of minor subcomplexes (except for Y complex) have been obtained^21,25^.

As the core scaffold of NPC, the inner ring (IR) attaches itself to the cytoplasmic, nucleoplasmic and luminal ring (CR, NR and LR), and is directly involved in the formation of the central transport channel of NPC. Here, using single-particle cryo-EM, we report the near atomic structures of IR, which reveals multiple interaction modes and assembly mechanism of IR.

## Results

### To obtain the structure of entire NPC by single particle analysis

Previous studies and our preliminary experiments showed great structural variability of NPC ^15,16,18,34^ (Supplementary information, Fig. S1a). To enrich high-quality samples of *sc*NPCs, we optimized and developed a gentle and rapid purification method referring to previous published literatures ^17,35^ to increase the mono-dispersity and uniformity. The key step is to treat cells with a chemical reagent-alpha-mating factor (α factor) (Supplementary information, Fig. S1b), which arrests cell in G1 phase synchronously and makes cells present a recognizable Shmoo shape (Supplementary information, Fig. S2) ^35^. To examine the impact of this medicine on structure of NPC, two subsets of data were collected. One was NPC treated with α factor, and the other one without α factor. Finally, we obtained two low resolution structures of NPC at 24 Å resolution (+ α factor, with 32159 particles) and 26 Å resolution (-α factor, with 21542 particles), respectively (Supplementary information, Fig. S3). Structural comparison indicated no obvious structural differences for +/− α factor treatment, especially in the region of IR (Supplementary information, Fig. S3). Because the particles on the grid displayed a certain preferential orientation (Supplementary information, Fig. S1), we collected data combining the tilting angles at 0 and 40 degrees. We acquired 279,900 particles from 283,881 and 12,939 micrographs at tilting angles of 0 and 40 degrees respectively, and determined the cryo-EM map of entire NPC at approximately 12.0 Å using final 51,220 particles by single particle analysis (SPA) (Supplementary information, Figs. S4a, S5, a-d) (refer to the Materials and methods for details). In agreement with previous reports ^17,18^, the resulting map of entire NPC presents a sandwiched architecture with one layer of IR and two layers of outer ring (CR and NR) and exhibits eightfold rotational symmetry. Besides, the controversial LR that surrounds the periphery of NPC, is visible in our map (Supplementary information, Fig. S5d). The LR was also found in tomography of *x*/NPC (*Xenopus laevis* NPC) ^22^ and detergent-extraction *sc*NPC ^17^, but not in tomography of *sc*NPC ^18^, which was probably due to the limited resolution or strong noise of nuclear envelope or its intrinsic flexibility. *sc*NPC spans 60 nm along the nucleocytoplasmic axis, and nearly spreads 100 nm parallelly to the equatorial plane of NPC (Supplementary information, Fig. S5d). When we gradually increased the map’s threshold, only IR region becomes clearly visible among the three layers (Supplementary information, Figs. S3b, S5d), indicating that IR is the most stable and uniform region in the entire *sc*NPC, which is consistent with the function of IR as the core scaffold. The IR measures 80 nm and 43 nm in outer and inner diameters, respectively, which is the same as that published by Seung JK *et al.* and narrower than its in-cell diameter (Fig. 1b; Supplementary information, Fig. S6a) ^17,18^. Although the diameters of entire IR vary even within the same species or across different cellular states ^17,19,36^, the asymmetric element of IR, here namely IR monomer, is very similar in structural characteristics even from different species (Supplementary information, Fig. S6b), suggesting that the IR monomer is relatively conserved and rigid in the variable IR.

**Fig. 1.**
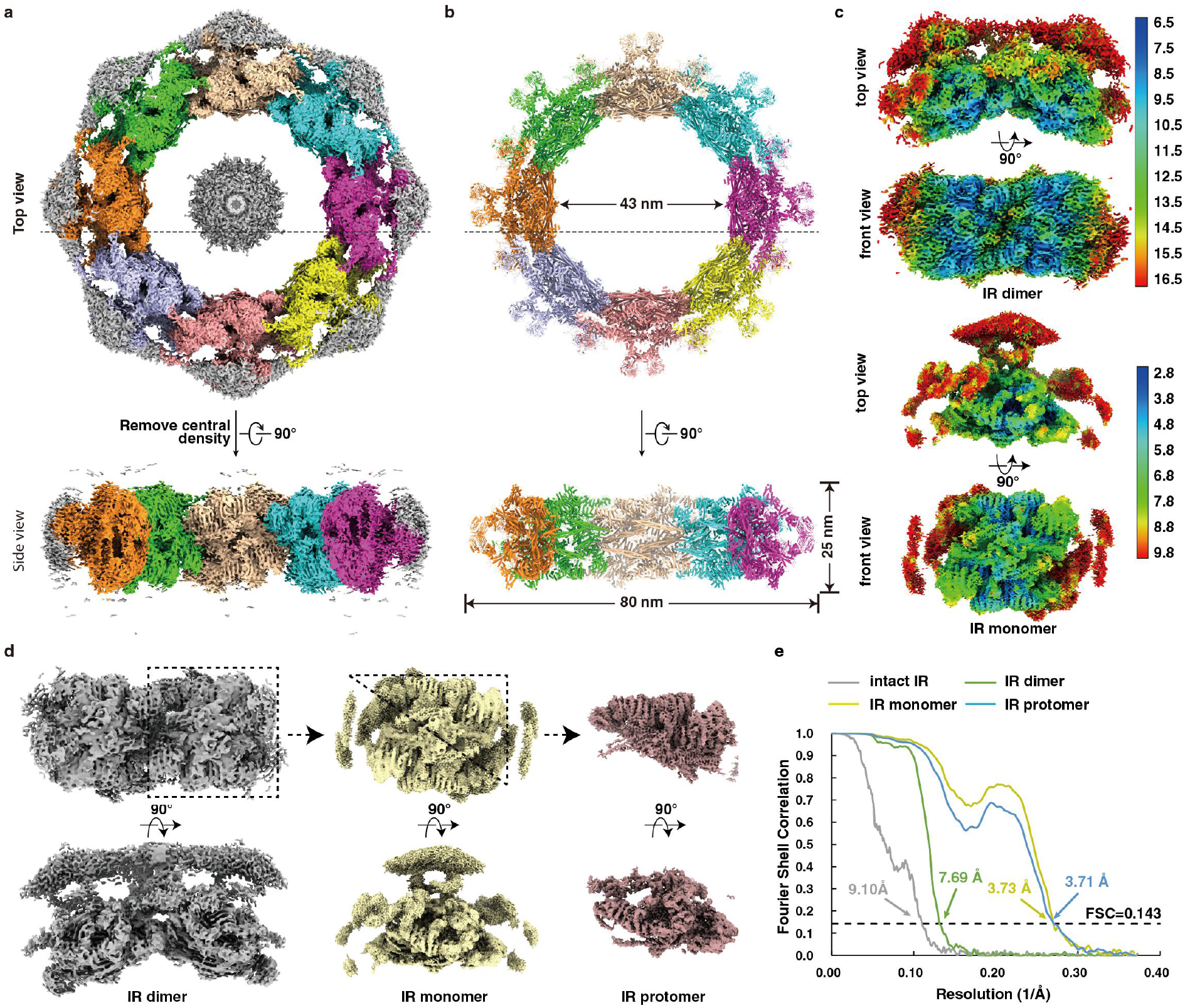
Cryo-EM structures of the inner ring (IR) of the *Saccharomyces cerevisiae* NPC. (**a**, **b**) Cryo-EM density map (a) and the atomic model (b) of the intact IR. The map and model are shown in top view (upper panel, dotted line indicated the cross section corresponding to the side view) and side view (lower panel). Different colors represent different IR monomers. Threshold contour level is 0.22. (**c**) Density maps of IR dimer, IR monomer and IR protomer determined in this study. (**d**) Color-coded 3D reconstructions of IR dimer (upper panel) and IR monomer (lower panel) showing local resolutions in different views. Threshold contour levels of dimer and monomer are 0.2 and 0.13, respectively. The local resolutions were estimated with cryoSPARC and generated in ChimeraX. (**e**) Gold standard Fourier shell correlation (FSC) curves for the maps of intact IR, IR dimer, IR monomer and IR protomer.

### To obtain structure of IR at 3.73 Å

Next, we tried to improve the resolution of nodical IR region from the entire NPC map. Firstly, we extracted one IR monomer of each NPC particle from raw micrographs based on the angular parameters of intact NPC and the relative position between IR monomer and NPC. And then we refined the CTF and angular parameters of the extracted IR monomers. Finally, the optimized parameters were applied back to the reconstruction of the entire IR (refer to the Materials and methods for details). By using this strategy, we obtained a final density map of entire IR at 9.10 Å resolution (Fig. 1, a, e; Supplementary information, Figs. S4b, S5e), which appears to be shaped like a “Finger Ring” (Fig. 1a). In order to further improve the resolution of IR monomer, we extracted each IR monomer and combined them for further processing (Supplementary information, Figs. S7, S8a). The overall resolution of the resulting IR monomer is 3.73 Å with a local resolution at 2.8-3.4 Å at the interaction region between Nup188 and Nup192 (Fig. 1c), where side chains are clearly visible for most residues (Supplementary information, Figs. S9a, S9b). In order to improve the resolution to aid model building, we performed symmetry expansion of each IR monomer particle, which resulted in twofold particles of IR protomer (Supplementary information, Fig. S8a) (refer to the Materials and methods for details). Although the final resolution of protomer is 3.71 Å (Fig. 1c; Supplementary information, Fig. S7e), only slightly higher than that of IR monomer, the map quality of CNT is much better than that in the IR monomer.

Since the quality is poor at the peripheral region of the IR monomer map, we then performed the reconstruction of the IR dimer to improve the map quality at the connection region between the IR monomers. A reconstruction of IR dimer was obtained at 7.69 Å resolution (Fig. 1, d, e), where Nup170 dimer and Nic96 molecules are clearly recognized (Fig. 1c; Supplementary information, Fig. S9c).

The much-improved quality of IR monomer enables us to place IR subunits into correct orientations more accurately than before. Special S-shaped Nup188 and Nup192 are easily identified in the map with local near-atomic resolution, and the representative long stalked α-helices from CNT complexes close to Nup188 and Nup192 are also conspicuous (Fig. 1c; Supplementary information, Fig. S7e). Many α-helices of α-solenoid domain from Nup170 and Nic96-B-1/2 are seamlessly fitted into the tubular densities that are characteristic of α-helices (Supplementary information, Fig. S9b). Although the peripheral density containing Nup157 and Nic96-A-1/2 exhibits poor quality, we combined with previous integrative models ^17,18^ to recognize them, and the final models of Nup157 and Nic96-A-1/2 fit well into the corresponding EM maps of IR monomer and IR dimer, with cross-correlation coefficients at 0.83 and 0.84 respectively.

We finally generated an atomic model of IR monomer, which is composed of two copies of large subunits, Nup188, Nup192, Nup157 and Nup170, and four copies of Nic96 and CNT complex (including Nup57, Nup49 and Nsp1) (Supplementary information, Figs. S10, S11a). Eight copies of IR monomer were docked into the above entire IR reconstruction to generate a near atomic resolution model of IR including 192 subunits and accounting for ~ 1/3 mass of entire NPC with approximately 16 MDa (Fig. 1b; Supplementary information, Figs. S10, S11a). Comparison with previously published integrated IR model from yeast (*18*), almost every IR subunit has different orientation (Supplementary information, Fig. S11b).

### To determine the structure of full-length Nup188 by purified protein

Nup188 and Nup192, the two of the largest subunits in NPC, are evolutionary conserved homologous nucleoporins. Since the density maps of Nup188 and Nup192 are not completely resolved in the IR monomer map, we next performed SPA for the purified Nup188 and Nup192 samples, respectively, and succeeded with the full-length structure of Nup188 at a resolution of 2.81 Å (Supplementary information, Fig. S12). Overall Nup188 appears to be a cray shape with a dimension of ~ 170 Å × 90 Å × 65 Å (Fig. 2b). The N-terminal contains two “crab claw” domains, calmp1 and clamp2, and between them is the predicted conserved SH3-like domain. The Neck domain is at the corner followed by a crescent C-terminal domain (CTD) (Fig. 2, a, b). The relatively poor density of the SH3-like domain and the tail of CTD represent intrinsic flexibility of these regions (Movies 1, 2), which is consistent with their roles in mediating interaction with neighboring IR monomer (Fig. 6f). The clamp2, Neck and CTD domains further form a large right-handed super-helical ring that resembles “S” letter including 52 stacked helices (Fig. 2c; Supplementary information, Fig. S13a). Surrounded by the axis of the “S”, two layers of concentric α-helices spiral up in parallel and are referred to the outer and inner layer with respect to their distances to the axis, which probably enables Nup188, like a spring, to stretch and shrink to a certain extent (Fig. 2c). The Nup188 map from SPA (Nup188_SPA_) fits into IR monomer map very well and has a good match with Nup188 map from IR monomer (Nup188_IR_), which shows that the obtained IR monomer map by SPA has high precision and accuracy (Supplementary information, Fig. S14a).

**Fig. 2.**
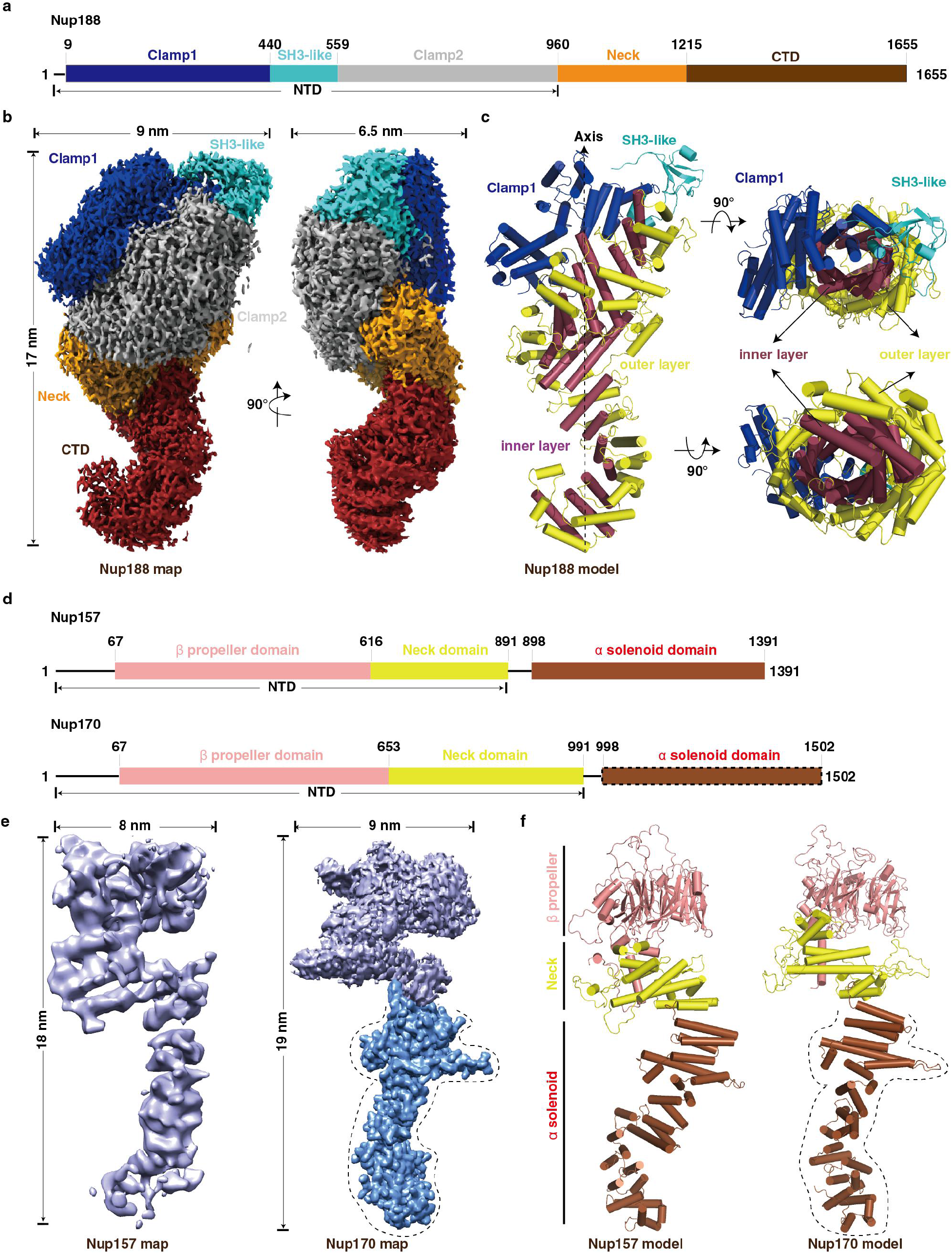
Structures of the full-length Nup188, Nup157 and Nup170 from purified proteins. (**a**) Schematic representation of the domain structures of Nup188. Domains are color coded. (**b**) Final cryo-EM map and diameter of Nup188. Color of domains are shown as in (a). (**c**) Model of Nup188. Colors of Clamp1 and SH3-like domains are shown as in (a), and S domain is divided into two layers-inner layer and outer layer, colored by raspberry and yellow, respectively. (**d**) Schematic representation of the domain structures of Nup157 and Nup170. Domains are color coded. α solenoid domain (boxed by dashed line) in Nup170 indicates missing regions in our map of Nup170. (**e**, **f**) Final cryo-EM maps (e) and models (f) of Nup157 and Nup170, respectively. Colors of domains and dotted line marked regions are shown as in (d). Cylinders represent α-helices. NTD, N-terminal domain; CTD, C-terminal domain.

Due to aggregation and a modest yield in both prokaryotic and eukaryotic expressing systems, we failed to resolve the structure of Nup192 by SPA. We generated the homologous model of Nup192 based on the structure of *ct*Nup192 (*Chaetomium thermophilum* Nup192), which is similar with the Nup188 model (Supplementary information, Figs. S13b, S14b). Compared with Nup188, Nup192 has an opening between clamp1 and clamp2 and a shorter tail in CTD. Another notable feature of Nup192 is the longer tower helix near the CTD domain, which is shorter in Nup188 and clearly visible in Nup192 map from IR monomer (Supplementary information, Fig. S14, b, c). Moreover, the distinguishing feature of Nup188 different from Nup192 is the existence of SH3-like domain in N-terminal, which is composed of several β-strands and loops (Fig. 2c and Supplementary information, Fig. S14c)^37^. The overall common S-shaped structural properties of Nup188 and Nup192 may play important roles in regulating and maintaining the malleability of IR.

### To determine the structures of full-length Nup157 and Nup170 by purified proteins

Nup157 and Nup170 are two homologous nucleoporins. Previous studies have reported that MBM motifs of Nup157 located at N-terminal are required for membrane binding and crucial for NPC assembly ^23,38^, suggesting that Nup157 and Nup170 might be helpful for anchoring of IR to NE. From the density maps of IR monomer and IR dimer (Fig. 1; Supplementary information, Fig. S15), we found that the N-terminals of Nup157 and Nup170 protrude into the nuclear envelope, indicating that the N-terminal domains of Nup157 and Nup170 participate in conjunction between IR and LR. To further gain insight into the structures of Nup157 and Nup170, we performed SPA for the purified Nup157 and Nup170 samples, respectively. The structures of the full-length Nup157 and the N-terminal of Nup170 were resolved with a resolution at 5.9 Å and 3.7 Å respectively (Supplementary information, Figs. S16, S13, c, d). Integrative models of the full-length Nup157 and Nup170 were produced by combining the SPA results with the previous x-ray structures of the N-terminal of Nup157 (PDB 4MHC) and the C-terminal of Nup170 (PDB 3I5P) and the maps of IR monomer and IR dimer obtained in this study. Nup157 and Nup170 are structurally spoon-shaped and composed of three distinct regions, a compact β-propeller domain followed by a short-stalked Neck domain and a long super-helical α solenoid domain. The β-propeller domain and the Neck domain further form a U-shaped N-terminal domain (Fig. 2, d-f). The full-length of Nup157 has dimensions of approximately 180 Å high and 80 Å wide. Structural analysis suggested that Nup157 is intrinsically flexible, especially in the α-solenoid domain (Supplementary information, Fig. S16g, Movie 3), which might hint its versatile role in regulating IR and LR. Different from the previously reported C-shaped negative stain structure ^39^, Nup170 shows a similar structure with Nup157, but with slightly larger dimensions, approximately 190 Å high and 90 Å wide (Fig. 2, e, f), which is reminiscent of its possible synergistic function with Nup157. Comparing the NTD of Nup157 and Nup170, one obvious difference is that there are two protruding horns at the bottom of “U” from Nup170 (Supplementary information, Fig. S17g). Easy degradability of Nup170 during purification suggests that its C-terminal is more flexible than that of Nup157 (Supplementary information, Fig. S17a). The flexibility of Nup157 and Nup170 might reflect their functional roles in assembly and variability of NPC.

### Extensive interactions within IR monomer

Based on the atomic model of IR monomer, we analyzed the interactions between the subunits. The IR monomer can be divided into three layers (Supplementary information, Fig. S10). The outer layer, shaped like a triangular rooftop from the top view, is comprised of two copies of Nup157 (Nup157-1/2), two copies of Nup170 (Nup170-1/2) and four copies of Nic96 (Nic96-A-1/2, Nic96-B-1/2) (Fig. 3a; Supplementary information, Fig. S10, Movie 4). The middle layer contains two copies of Nup188 and two copies of Nup192, which assembles into a pore-facing “Z”-shaped pattern with an arched cavity on the cross section. The inner layer includes four CNT complexes (CNT-A-1/2, CNT-B-1/2) and each CNT complex composes of three homologous proteins, Nup57, Nup49 and Nsp1 (Supplementary information, Fig. S10, Movie 4). In our model, the four CNT complexes assemble into a rhombic tetramer residing in the arched cavity of the middle layer (Fig. 3E and Supplementary information, Fig. S10), which is directly involved in the formation of the central transport channel of NPC (Fig. 1b; Supplementary information, Fig. S10).

**Fig. 3.**
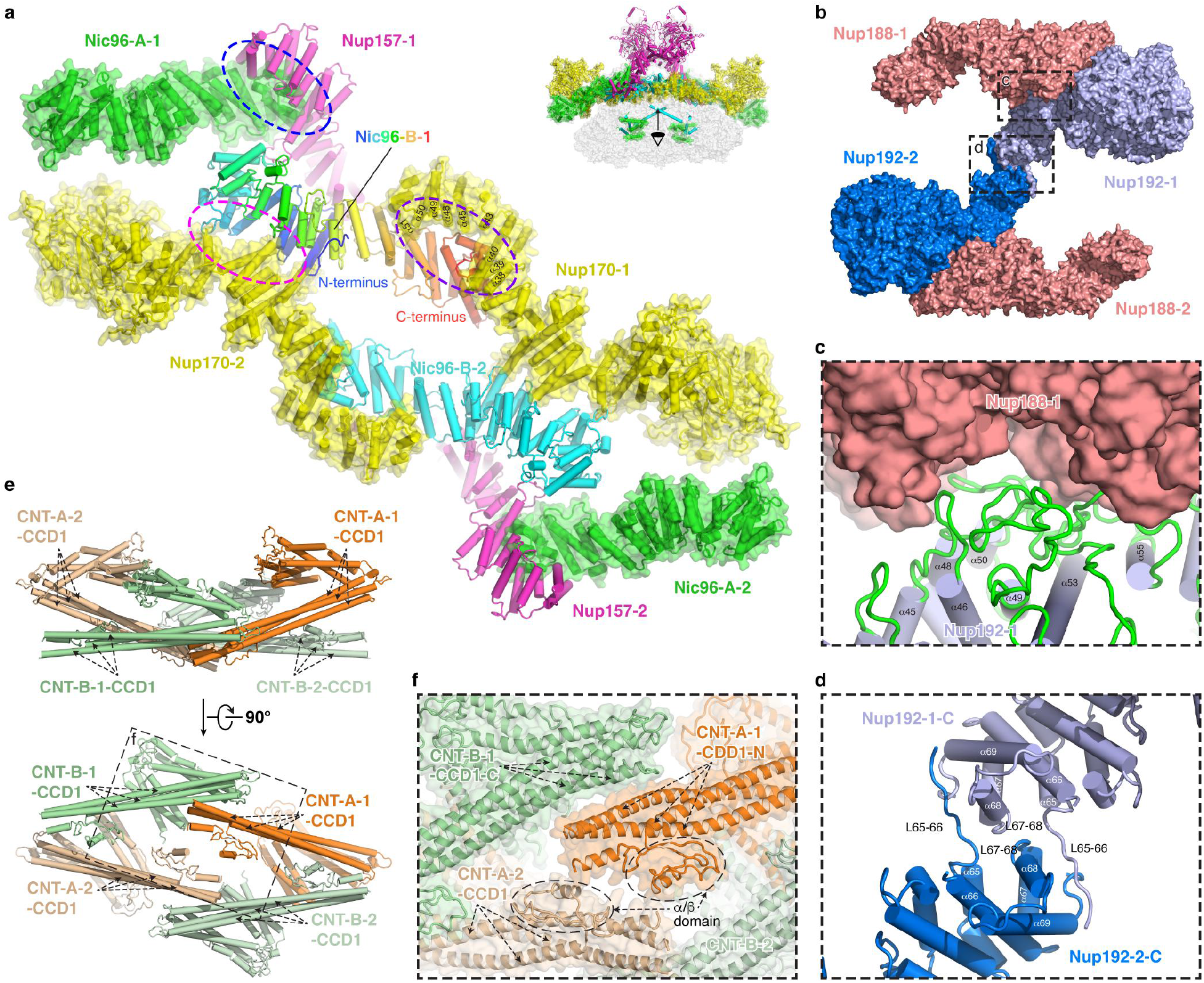
Intra-layer interactions in the IR monomer. (**a**) Interactions within the outer layer. Purple, red and blue dashed ellipses represent interactions between Nic96-B-1 and Nup170-1, Nic96-B-1 and Nup170-2, and Nic96-A-1 and Nup57-1, respectively. (**b**) Interactions within the middle layer. Two interfaces are boxed by black dashed line and enlarged in (c) and (d), respectively. (**c**) Details of interaction between Nup188-1 and Nup192-1. (**d**) Details of interaction between two Nup192 proteins. (**e**) Interactions within the inner layer. The black dashed box is zoomed in (f). (**f**) Details of the interaction between CNT complexes.

Within the outer layer, N-terminals of Nup157-1/2 and Nup170-1/2 extend into the LR (Supplementary information, Figs. S10, S15). The C-terminals of Nup170-1/2, together with four diagonally arranged Nic96, form a slightly curved plane facing towards the pore of NPC, just like a bridge deck. Nic96 plays a key role in this assembly. Its C-terminus is embedded in a groove formed by the helices of α38-α40, α43, α45 and α48-α51 from the C-terminal α-solenoid domain of Nup170 (Fig. 3a, purple circle; Supplementary information, Fig. S13, d, e). Meanwhile, its N-terminus directly contacts with the N-terminal region of α-solenoid domain of the other Nup170 via α-helices and loops (Fig. 3a, magenta circle). By this way, two Nic96 molecules (Nic96-B-1/2) and two Nup170 molecules (Nup170-1/2) are linked together and form a rhomboidal plane, which is the center of the bridge deck. Nup157-1/2 is anchored to this bridge by association of its C-terminus with the C-terminus of Nic96-A-1/2 (Fig. 3a, blue circle; Supplementary information, Fig. S13, c, e), consistent with the results of cross-linked mass spectrometry ^17^.

For the middle layer, benefited from the full-length structure of Nup188 and the improved resolution of IR monomer, we found an extensive compact interface between Nup188 and Nup192, with ~ 2157 Å^2^buried surface area (Fig. 3, b, c), proving the two large subunits’ direct interaction, which is distinct from the *in vitro* results ^40^. The interaction interface is mainly composed of flexible loops and small α-helices between residues ~ 174-819 of Nup188 and residues ~ 1060-1432 of Nup192 (Fig. 3c; Supplementary information, Fig. S13, a, b). Moreover, two question-marker Nup192 molecules dimerize as arched shape with ~ 968 Å^2^ interface area via interactions between the flexible C-terminal α-helices α65-α69 and loops L65-66 and L67-68 from each other (Fig. 3d; Supplementary information, Fig. S13b).

The three homologous proteins forming CNT complex in the inner layer are structurally featured by the canonical heterotrimeric coiled-coil domains (CCD1, 2 and 3) for all proteins and an additional α/β domain for Nup57 ^28^ (Supplementary information, Fig. S13, f-h). Linkages between four CNT heterotrimers are primarily through the CCD1. In detail, the N-terminal of CCD1 in CNT-A-1 is adjacent to the C-terminal of CCD1 in CNT-B-1, and simultaneously, the tips of CCD1 including the α/β domain from CNT-A-1 and CNT-A-2 are closely associated (Fig. 3, e, f). In the rhombic tetramer each CNT monomer contributes a long cylindrical CCD1 domain, which is situated in the innermost part of the inner layer (Supplementary information, Fig. S10). The N-terminals of CCD1 have been identified as the place where the extremely flexible FG repeats locate ^41,42^. The lack of the FG repeat motifs in the model is likely due to their intrinsically disordered properties. The CNT rhombic tetramer has an inside room, which may provide a space for elastic deformation of the CNT complexes (Fig. 3e; Supplementary information, Fig. S10).

A lot of interactions between different layers exist within the IR monomer (Fig. 4a). Nup170 of the outer layer makes extensive interactions with the middle layer (Fig. 4, a-e). The C-terminal of Nup170 sits on the concave surface at the junction of Nup188 and Nup192 (Fig. 4, b, c) and interacts with both proteins (Fig. 4, d, e). It binds to Nup188 by association of its α-helices α39, α41 and α43 with the α-helices α27, α29, α32 and two long loops L26-27 and L29-30 from N-terminal of Nup188 (Fig. 4d). It also interacts with Nup192 through the binding of its α-helices α38-α43 and loops L38-39, L40-41, L42-43 and the C-terminal loop with α-helices α58, α59, α61, as well as loops L56-57, L58-59, and L60-61 from Nup192 (Fig. 4e). Moreover, agreement with the biochemically identified interaction between Nup188 and Nic96 ^17^, the α-helix α13 and following loops L14-15 and L15-16 in the N-terminal of Nic96-B-1 connects to α-helices α60, α62, α63, α66 and α57 in the C-terminal of Nup188 (Fig. 4f). These interactions contribute to the conjunction between the outer and middle layers.

**Fig. 4.**
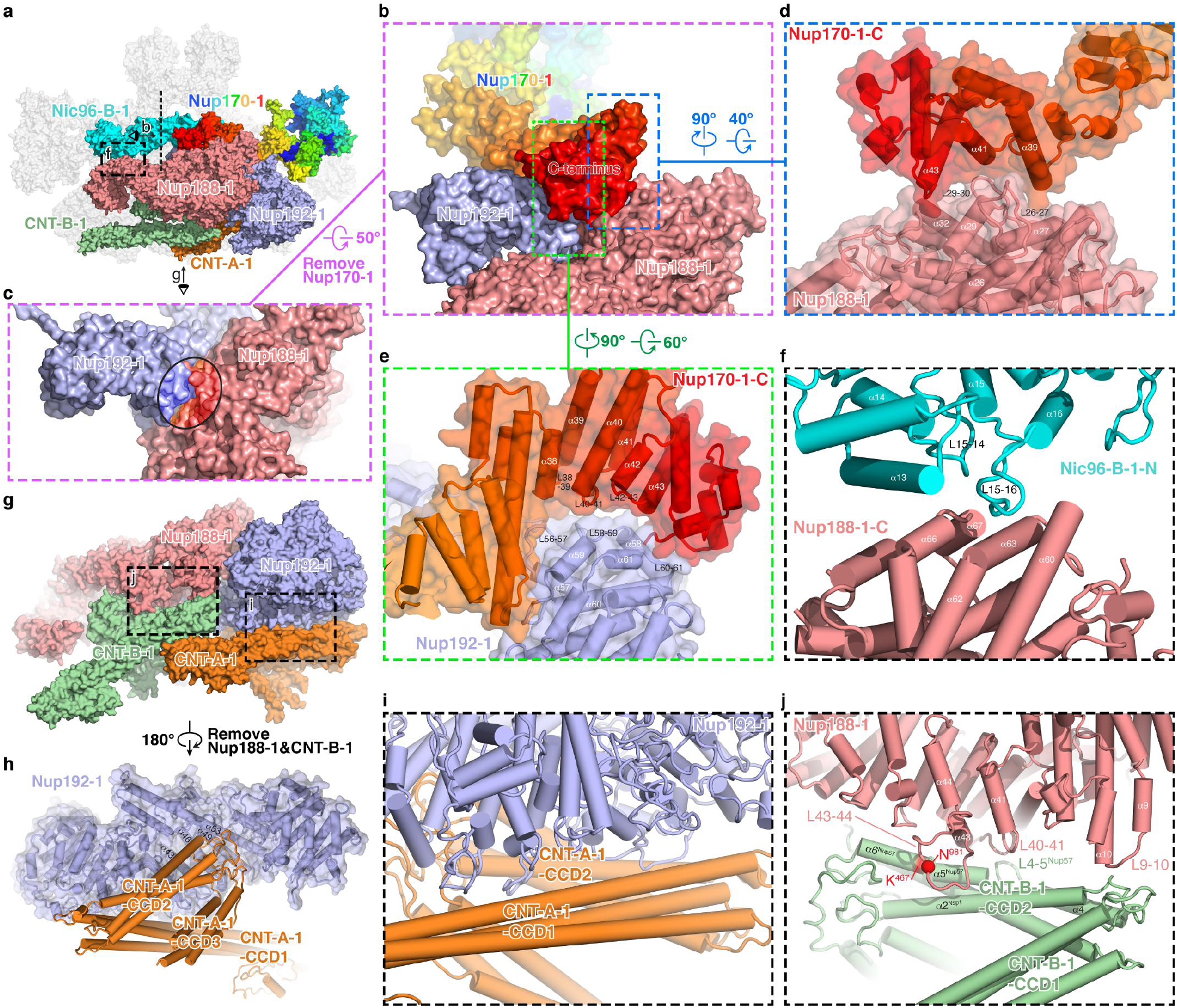
Inter-layer interactions in the IR monomer. (**a**) Overall view of subunits involved in the inter-layer interactions, with eye symbols and arrowheads indicating view directions shown in the following panels. (**b**) Interactions of Nup170 with Nup188 and Nup192. Two interfaces are boxed by blue and yellowgreen dashed lines and enlarged in (d) and (e), respectively. (**c**) The concave surface at the junction of Nup188 and Nup192. The area contacting to Nup170 is circled. (**d**, **e**) Details of interaction of the C-terminal of Nup170-1 with Nup188-1 (d) and Nup192-1 (e). (**f**) Details of interaction between Nic96-B-1 and Nup188-1. (**g**) Overall view of subunits involved in the middle and inner layer interactions. Two interfaces are boxed by black dashed lines and enlarged in (i) and (j), respectively. (**h**) Loops connecting CCD2 and CDD3 of CNT-A-1 protrude into the groove located in the “S” domain from Nup192-1. (**i**) Long stalked CCD1 and CCD2 of CNT-A-1 clamp the N-terminal corner of Nup192-1 by extensive contacts via α-helices and loops. (**j**) Details of interaction between Nup188-1 and CNT-B-1.

Connections between the middle and inner layers are mediated by the interaction of Nup188 and Nup192 with CNT complexes (Fig. 4g). For the interaction between Nup192 and CNT, the loops connecting CCD2 and CDD3 of CNT-A-1 protrude into the groove formed by α43, α46, α49 and α53 located in the “S” domain from Nup192-1 (Fig. 4h), and block the hole formed by the tower helix and C-terminal domain of Nup192 (Supplementary information, Fig. S18). In addition, the long stalked CCD1 and CCD2 of CNT-A-1 clamp the N-terminal corner of Nup192-1 by extensive contacts via α-helices and loops (Fig. 4i). Two interaction interfaces exist between Nup188-1 and CNT-B-1 (Fig. 4j). One interface is contributed by the CNT surface formed by α-helix α5 and loop L4-5 from Nup57, and α-helix α2 from Nsp1, and the Nup188 surface formed by α-helices α41 and α43, as well as loops L40-41 and L43-44 (Fig. 4j). This interaction has been verified by a pair of cross-linked residues identified from the endogenous *sc*NPC, Asn^981^ from Nup188 and Lys^467^ from Nup57 ^17^. The other interface is formed by the loops linking CCD1 and CCD2 of CNT-B-1, and α-helix α10 and loop L9-10 of the N-terminal Nup188 (Fig. 4j). The above multiple interactions between CNT and Nup188/192 may provide ways for the shift of conformational changes from the inner layer CNTs to the middle layer subunits.

In addition, several unassigned densities were found in the C-terminals of Nup188 and Nup192 and the surrounding of the CCD3 of CNT, most likely as part of the flexible N-terminal of Nic96 based on the crystal structure of Nic96-CNT, biochemical results and the predicted structure of the full-length Nic96 (28,38,42,46) (Supplementary information, Fig. S19). In this case, four Nic96 molecules string all three layers of IR monomer and interact with all IR subunits, suggesting its indispensable role as the keystone of NPC assembly (Supplementary information, Fig. S19). Moreover, located between Nic96-A-1 and the N-terminal of Nup170, an unambiguous density might come from the Nup53/Nup59 based on the mass spectrometry and biochemical results (Supplementary information, Fig. S9c) ^17,36^, indicating the possible function of the two flexible linker nucleoporins in assembly of IR.

### Elastic CNT tetramer in the center of transport channel

As mentioned above, four CNT complexes are arranged as a rhombic tetramer, which connects one by one to form the innermost circle of IR (Supplementary information, Figs. 3e, 1, a, b). Thus, two approximately parallel rhomboidal structures of the inner and outer layers, together with the Z-shaped Nup188-Nup192 middle layer, constitute a sandwiched construction of IR monomer (Supplementary information, Fig. S10, Movie 4). The structure of the whole IR is not very compact as demonstrated by the presence of a lot of unfilled spaces in it (Fig. 1, a, b). From a mechanical point of view, above special multilayer sandwiched assembly and the relatively loose structure might lead to an elastic mechanism, which could mediate the transfer of the conformational changes from the inner layer to the middle and outer layers. The CNT complex in the inner layer (CNT monomer) contains three cylindrical CCD domains connected by two hinges (H1 and H2) (Fig. 5). The hinge region is unstable and prone to change. So, total eight hinge regions in the CNT tetramer could allow the entire tetramer to stretch horizontally and/or vertically (Fig. 5), which make the CNT tetramer as an elastic element. The innermost elements of CNT are the long cylindrical CCD1 domains, where FG repeats form the diffusion barrier of the central transport channel ^41,42^. The cargo may be sensed by FG repeats, which may further induce conformational changes of CCD1 domains. Therefore, it is reasonable to deduce that the structural variability of the CNT tetramer could be an adaptation to the passage of cargoes of different sizes by own structural change.

**Fig. 5.**
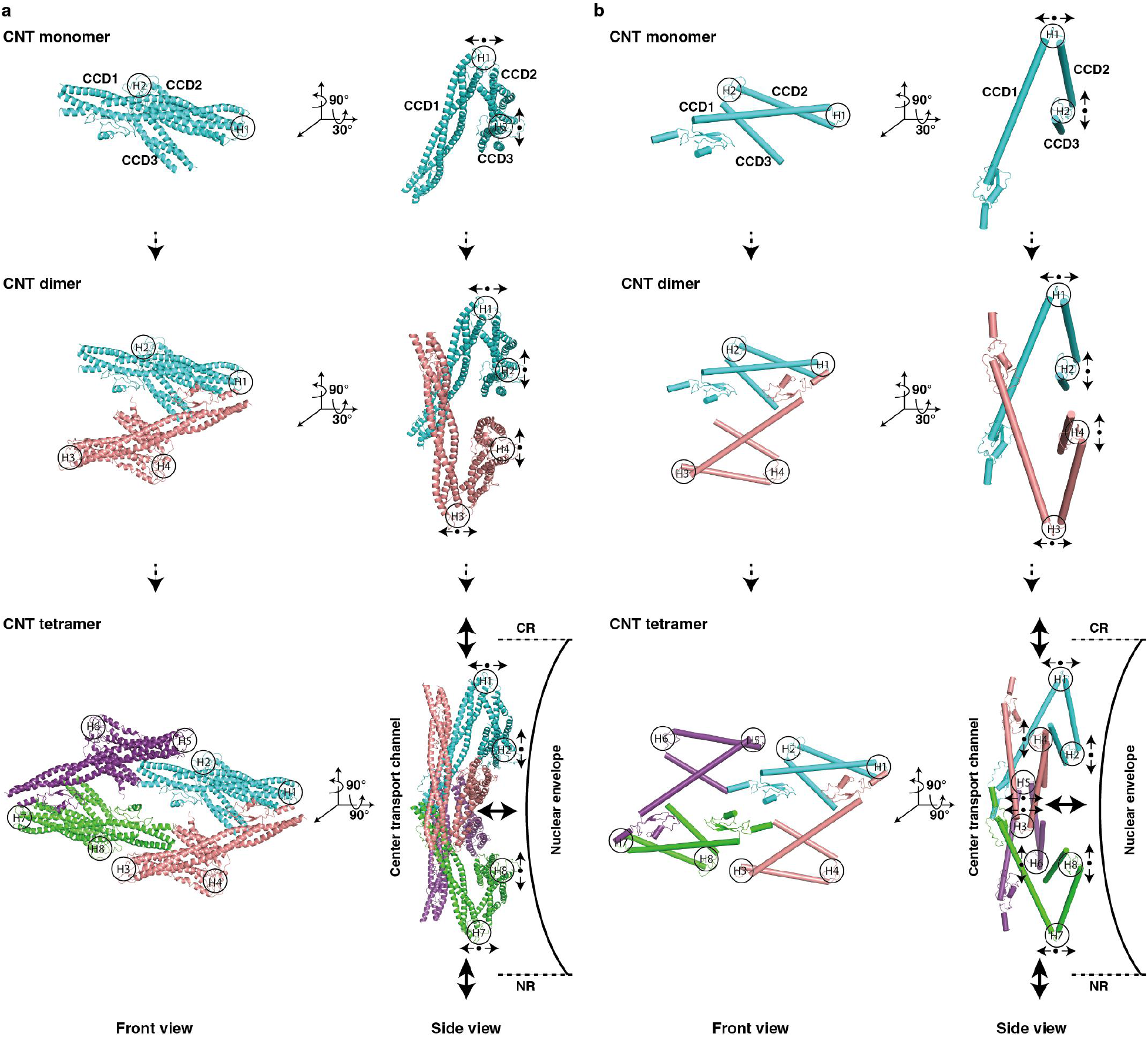
Conformational change of CNT complex. Molecular architectures and model illustrations are shown in (**a**) and (**b**), respectively. Top and side views are shown in upper and bottom panel, respectively. Among three CCD domains in CNT monomer, two hinge regions (H1 and H2) regulate angular variation of adjacent CCD domains to sense cargo with different sizes. Small black arrows represent possible motion of each CCD domain around hinge region. The resultant motion of CNT tetramer is shown in larger black arrows.

### Molecular mechanism of dilation and constriction of NPC

Comparison of the IR monomers in various states and from different species indicates the great similarity in composition, structure and size (Supplementary information, Fig. S6). Our cryo-EM structure of IR reveals that, as described in the above sections, three core subunits-Nup188, Nup192 and Nic96 in each IR monomer have extensive interactions with all subunits of the IR monomer (Fig. 4 and Supplementary information, Fig. S19), forming a connection network to maintain the IR monomer in stable and elastic structural state. Comparing with this, the interactions between IR monomers are rather different. We found that the inter-monomer interactions have loose and instable properties. In the outer layer, two Nup170 molecules from the neighboring IR monomers form a herringbone conformation through the contacts of N-terminals via extensive interactions through loops (Fig. 6a). Besides, we also found a poor density region linking the SH3-like domain of Nup188-1/M1 and the N-terminal of the adjacent Nic96-A-1/M2 (Fig. 6f). In the middle layer and inner layer, the α-helix α17 and loop L17-18 of the N-terminal of Nup192-1/M1 have small interaction with the α-helices α72-α73 and loop L72-73 from the flexible C-terminal of the neighboring Nup188-1/M2 (Fig. 6c). In addition, the long loop L23-24 from Nup192-1/M1 interacts with the loops connecting CCD1 and CCD2 of the neighboring CNT-A-2/M2 (Fig. 6d). Moreover, a binding region is formed by the hinges between CCD1 and CCD2 from CNT-A-1/M1 and the neighboring CNT-A-2/M2 (Fig. 6e). All above interactions are weak and loose. Thus, our structure exhibits an interaction pattern of IR: relatively stable and strong interactions within IR monomer and relatively instable and loose interactions between IR monomers.

**Fig. 6.**
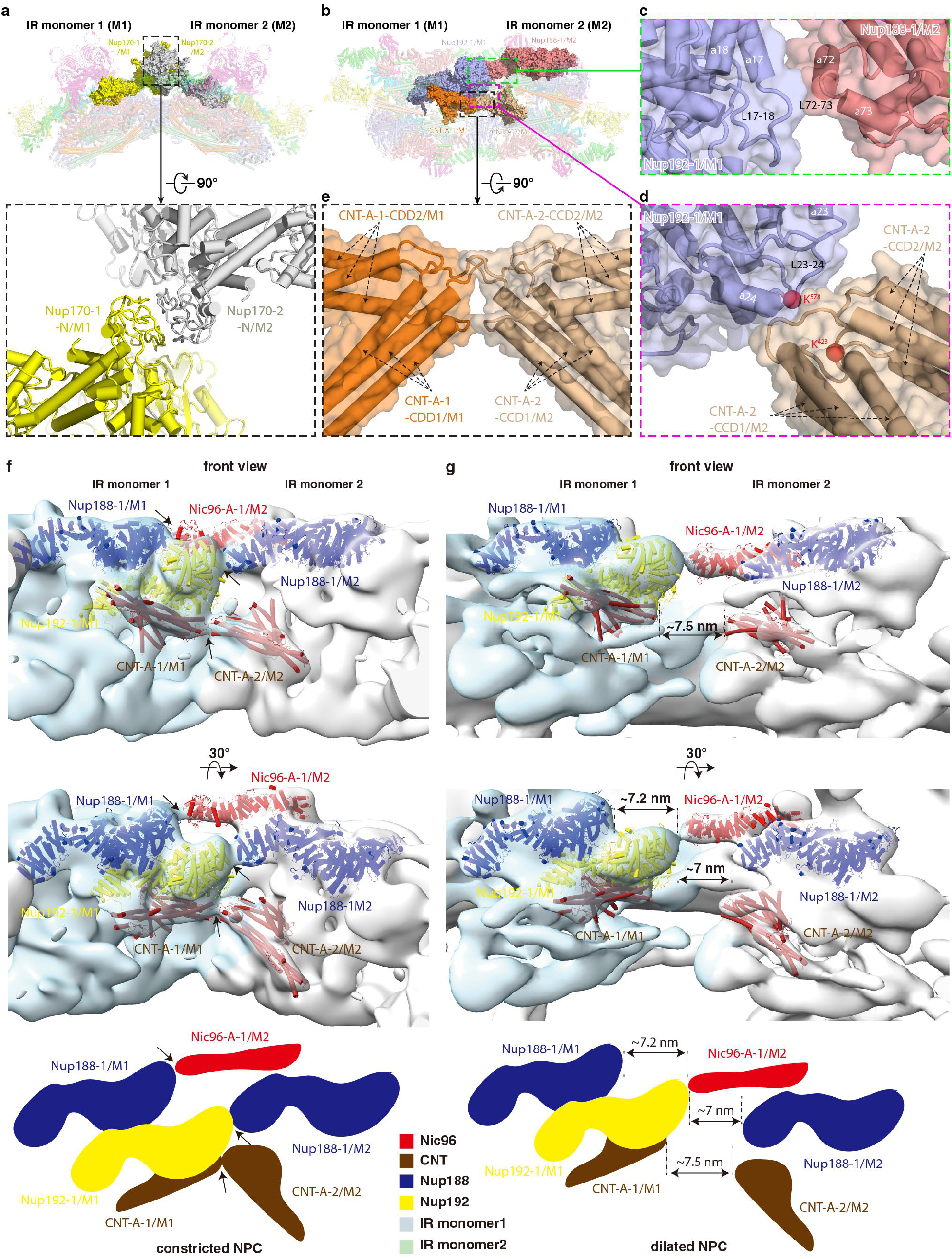
Interactions between adjacent IR monomers and comparison of dilated and constricted NPC. (**a**) Interaction between two Nup170 molecules from adjacent IR monomers. (**b**) Overall view of subunits from middle and inner layers involved in the inter-monomer interaction. Three interfaces are boxed by green, magenta and black dashed lines and enlarged in (c), (d) and (e), respectively. (**c**) Details of interaction between Nup192-1/M1 and neighboring Nup188-1/M2. (**d**) Details of interaction between Nup192-1/M1 and neighboring CNT-A-2/M2. (**e**) Details of interaction between CNT-A-1/M1 and neighboring CNT-A-2/M2. Distance change between IR monomers from constricted NPC (**f**) to dilated NPC (**g**). Two views are shown and schematic diagrams are drawn at the bottom. The representative subunits of IR are color coded. The interacting subunits of each interaction in the constricted NPC (black arrows) are separated by approximately 7 nm in the dilated NPC (dashed lines).

When we compared our IR structure with previously published integrated *in situ* yeast IR model ^18^, the IR diameter is significantly different, 80 nm in this study and 100 nm in the *in situ* study (Supplementary information, Fig. S6a). Close inspection found that the diameter difference primarily comes from the distance changes between IR monomers. In our structure, the disordered loop of N-terminal of Nic96-A-1 interacts the flexible SH3-like domain of Nup188-1 (Fig. 6f), and the weak density suggests this interaction is rather loose. While in the *in situ* study, Nic96-A-1 is switched to contact with the adjacent N192-1 and the aforementioned other interactions found in this study are separated with each other by about 7 nm (Fig. 6, f, g). As a result, about 90% perimeter change of IR is due to the different interaction modes between IR monomers and the resulting different distance between IR monomers. Thus, the NPCs observed in this study and in the *in situ* study likely present the NPCs in a constricted state and in a certain dilated state, respectively. In constricted state Nic96 interacts with neighboring Nup188 while in dilated state Nic96 changes to interact with neighboring Nup192. Together with the interactions between four Nic96 molecules and CNT tetramers in IR monomer, we propose the flexible N-terminal fragment of Nic96 therefore may function as a switch to regulate NPC the movement of NPC between constriction and dilation to adapt for the passage of various cargoes.

In summary, our cryo-EM structures describe in detail the hierarchical assembly of the *sc*NPC IR, which consists of approximately 192 copies from 8 Nups with an eight-folded rotational symmetry. Each IR monomer, as the unit of the entire IR assembly, forms a multilayer sandwich structure. Structural analysis reveals various interaction modes and extensive flexible connections within and between IR monomers, providing a structural basis for the stability and malleability of IR, and a molecular mechanism of the conformational change of NPC between constriction and dilation.

## Acknowledgments

We are grateful to X.Y. and D.S.L for model building. We thank X.M.L. and L.P.C for helpful discussion. We sincerely thank the staff at both the cryo-EM center of South University of Science and Technology and the Tsinghua University Branch of the National Protein Science Facility (Beijing) for their technical support on the Cryo-EM and High-Performance Computation platforms.

## Funding

The National Basic Research Program 2017YFA0504601 (S-FS)

The National Basic Research Program 2016YFA0501101 (S-FS)

The National Natural Science Foundation of China 31861143048 (S-FS)

The National Natural Science Foundation of China 32071192 (S-FS)

The National Natural Science Foundation of China 91954118 (SS)

## Author contributions

S.-F.S. supervised the project; Z.L. prepared the samples, performed the biochemical and yeast growth analyses; L.Z., S.C., G.H., X.P. and Z.L. collected the EM data and performed the EM analysis; Z.L. and L.Z., G.H., and S.C. performed the model building and the structure refinement; P.W. participated in the EM data collection; Z.L., S.S. and S.-F.S. analyzed the structure; Z.L., L.Z., G.H and S.C. jointly wrote the initial draft; S.S and S.-F.S. edited the manuscript.

## Data and materials availability

The EM density map of the intact NPC (EMDB: EMD-XXXXX), the intact IR (EMDB: EMD-XXXXX), the IR dimer (EMDB: EMD-XXXXX), the IR monomer (EMDB: EMD-XXXXX), the IR protomer (EMDB: EMD-XXXXX), Nup157 (EMDB: EMD-XXXXX), Nup170 (EMDB: EMD-XXXXX) and Nup188 (EMDB: EMD-XXXXX) have been deposited in the Electron Microscopy Data Bank (www.ebi.ac.uk/pdbe/emdb/). Atomic coordinates of the IR monomer (PDB: XXXX) and Nup188 (PDB: XXXX) have been deposited in the Protein Data Bank (www.rcsb.org). All other data and materials are available from the corresponding authors upon reasonable request.

## Supplementary Materials

Materials and methods

Supplementary information, Figs. S1 to S19

Tables S1 to S2

References (*43–63*)

Movies S1 to S4

