## Supplementary Materials for "Near atomic structure of the inner ring of the *Saccharomyces cerevisiae* nuclear pore complex"

**This PDF file includes:**

Materials and methods
Figs. S1 to S19
Tables S1 to S2
Captions for Movies S1 to S4

### Materials and Methods

#### Yeast strains and plasmids

The budding yeast *S. cerevisiae* strains and plasmids used in this study are listed in Supplementary Table 1. All plasmids were checked by sequencing.

#### Purification of the native *S. cerevisiae* NPC

The yeast strain *Saccharomyces cerevisiae* (W303a and W303α) was used throughout the procedure. In the *Mlp1-PrA/Nup84-3FH* strain, endogenous *Mlp1* and *Nup84* were tagged with PrA preceded by TEV protease cleavage sequence (ENLYFQG) and 3x FLAG plus 10x His sequence, respectively, by homologous recombination of polymerase chain reaction (PCR)-amplified cassettes. There was no significant difference in cell growth and morphology of wild type and Mlp1-PrA/Nup84-3FH yeast cells (Supplementary information, Fig. S2). To gain the high quality endogenous *sc*NPC, we developed a purification protocol based on previous published methods [17,43-46]. Briefly, yeast cells were grown at 30°C in yeast extract peptone dextrose (YPD) medium until reaching the middle log phase (OD600 0.8-1.2) and then treated with 10 μg/ml of the alpha-mating factor for 2 h to synchronize cells in the G1 phase prior to harvest by centrifugation. The pellets of yeast cells were resuspended in lysis buffer (20 mM HEPES-KOH pH 7.4, 20 mM NaCl, 2 mM $MgCl_2$, 50 mM potassium acetate, 0.1% (w/v) Tween-20, 0.5% (w/v) Triton X-100, 10% (v/v) glycerol) at a weight/volume ratio of 1:1, and then frozen in liquid nitrogen followed by cryogenically ground in a mill (ATS Scientific: Freezer/Mill 6875). Frozen cell powder was resuspended in the lysis buffer with 1/100 (w/v) protease inhibitor cocktail (Roche), and gently stirred for 2h at 4°C. The lysate was centrifugated by 2500 RCF for 5 min followed by 12000 RCF for 5min. The supernatant was collected and incubated with anti-FLAG beads (MERCK, A2220) for 2h at 4°C. After extensive wash by lysis buffer, the crude NPCs were eluted by 3x FLAG peptide (sigma-Aldrich: F4799) in the elution buffer (20 mM HEPES-KOH pH 7.4, 20 mM NaCl, 2 mM $MgCl_2$, 50 mM potassium acetate, 0.1% (w/v) Tween-20, 10% (v/v) glycerol). The eluent was further purified by size exclusive chromatography using a Sephacryl S-500 HR column (GE healthcare: HiPrep™ 16/60 Sephacryl S-500 HR) in the elution buffer. The fractions containing NPCs were pooled together and incubated with the Dynabeads coupled to rabbit IgG antibodies (sigma-Aldrich: I5006-10 mg) for 1h at 4°C. IgG Dynabeads were prepared using the Dynabeads antibody coupling kit (ThermoFisher: 14311D). After washing with elution buffer, NPCs were released by TEV protease (in-house) cleavage for 2h at 4°C in the desired volume of elution buffer. Finally, a magnet was applied to remove the beads and the supernatant containing the *sc*NPC was collected by centrifuging at 20,000 g for 10 mins.

#### Purification of the Nup188, Nup157 and Nup170

DNA fragments of IR subunits were amplified by PCR using *S. cerevisiae* cDNA as template and cloned into corresponding vector for eukaryotic and prokaryotic protein expression. Details of expression constructs are shown in Supplementary information, Table S1. After screening, we focused on purifications of Nup188, Nup157 and Nup170 that displayed the better performances. All subunits were purified with the same following procedure. The plasmids expressing the C-terminal Flag-tagged protein were transformed into *S. cerevisiae* (W303a) cells using the traditional lithium acetate protocol [47]. Single colonies were picked up and inoculated in 5 ml Synthetic Dextrose (SD) medium containing 100 μg/ml ampicillin and 50 μg/ml Kanamycin and grown overnight at 30 °C as the starter culture. In the following morning, the starter culture was added into 1 L of SD medium containing antibiotics for further culture. Cells were grown at 30 °C until the OD600 reached ~2.0-2.5, and then protein production was induced by adding 2% galactose for extra 4-6 h. The yeast cells were harvested by centrifugation at 4800 rpm/min for 10 mins, resuspended in lysis buffer (20 mM HEPES-KOH pH 7.4, 50 mM potassium acetate, 500 mM NaCl, 2 mM $MgCl_2$, 0.1% NP40, 5% (v/v) glycerol, 1/200 (v/v) protease inhibitor cocktail) at a weight/volume ratio of 4:1, frozen in liquid nitrogen and cryogenically ground in the mill. The cell powder was thawed using 2x volumes of lysis buffer and stirred for 1 h. The lysate was cleared by centrifugation at 18000 rpm/min for 30 mins. The supernatant was incubated with anti-FLAG beads for 2h at 4°C. Then the beads were washed extensively by lysis buffer and elution buffer (20 mM HEPES-KOH pH 7.4, 50 mM potassium acetate, 150 mM NaCl, 2 mM $MgCl_2$, 0.01% NP40, 5% (v/v) glycerol) in turn. Proteins were eluted by 200 μg/ml 3x FLAG peptide in the elution buffer. The eluent was applied to a Superose S6 3.2/60 column (GE Healthcare) with running buffer containing 20 mM HEPES-KOH, pH 7.4, 50 mM potassium acetate, 150 mM NaCl and 2 mM $MgCl_2$. Peak factions were collected and concentrated for cryo-EM sample preparation.

**Synchronization of Yeast cell**

Alpha-mating factor is a well-known pheromone secreted by mating type alpha (MATα) cells, and can arrest mating type a (MATa) cells in the G1 phase with a clear hallmark of “Shmoo” shape [35]. To find the proper concentration of alpha-mating factor and the time point of cellular arrest, we conducted synchronization experiments of wild-type and Mlp1-PrA/Nup84-3FH yeast cell under different conditions. Briefly, over-night cultured yeast cells were diluted to OD600 of ~ 0.2 followed by continuous shaking for 4-6 h at 30°C until cells reach the log phase (OD600 ~ 0.8). And then the cells were divided into three equal parts, which were added with 5 μg/ml 10 μg/ml and 15 μg/ml of alpha-mating factor, respectively. We monitored the cell cycle arrest by laser scanning confocal microscopy (Olympus FV1200) every hour (Supplementary information, Fig. S2). Finally, we chose 10 μg/ml of alpha-mating factor to treat cells for 2 h, because most cells present “Shmoo” shapes at this condition, where the concentration of the drug is lower, which means less toxicity.

**Cryo-EM sample preparation and single-particle data sets acquisition**

For NPC complex, the purified samples with 10% glycerol were checked by negative staining using a Tecnai Spirit at 120 kV (Thermo Fisher Scientific). The most homogenous samples were selected to prepare cryo-EM samples. 200 mesh Au-lacey carbon grids with continuous carbon support film (Electron Microscopy Sciences) were glow discharged in air, and each grid was mounted on forceps in a Mark IV Vitrobot (Thermo Fisher Scientific) at 4 °C and 100% humidity. 4 μL of sample drops were floated on the grid for 60 s and then about 3 μL of them were removed by absorbing using pipettor. Then, 4 μL of elution buffer without glycerol was added to the grid and the grid was plunge-frozen in liquid ethane after blotting. For nucleoporins (Nup188, Nup157 and Nup170), 4 μL of each purified sample was applied onto a plasma-cleaned holey carbon Au grid (Quantifoil, R1.2/1.3, 300 mesh). Grids were blotted at 100% humidity and 4 °C and then plunge-frozen in liquid ethane using a Mark IV Vitrobot.

Data sets of both NPC complex and nucleoporins (Nup188, Nup157 and Nup170) were collected on a Titan Krios electron microscope operating at 300 kV, equipped with a Gatan K3 Summit direct electron detector and a GIF Quantum energy filter. All data sets were collected using SerialEM [48] with same imaging settings. Images were recorded using a pixel size of 0.668 Å at a magnification of 130,000. The defocus value of each image was set from -1.5 to -2.0 μm during data collection. Each micrograph was dose-fractioned into 32 frames with a total dose of about 50 $e^-$/Å$^2$.

Finally, 296820, 16527, 15880 and 8451 movies were collected for NPC, Nup188, Nup157 and Nup170, respectively. The beam-induced motion of the whole micrograph with 32 movie frames was corrected by MotionCor2 [49]. CTF parameters were estimated by Patch CTF estimation in cryoSPARC [50] for NPC, Nup157 and Nup170, and were estimated by CTFFIND4 [51] for Nup188. Further details of data collection are given in Supplementary Table 2.

**Initial defocus estimation of NPC**

To obtain reliable defocus of particles in the micrographs, we used CTFFIND4, Gctf [52] and patch CTF estimation in cryoSPARC to estimate the CTF parameters. The CTF parameters from different software didn't make much difference. However, we found that for bad micrographs that contained crystalline ice, contamination or carbon films, the CTF parameters from different software were much different. According to the estimations calculated by patch CTF estimation, 187,076 micrographs were selected from 296,820 micrographs with the maximum resolution value below 4 Å, the astigmatism value below 1000 Å and the defocus ranged from 0.4 μm to 4 μm. Considering that the diameter along the equatorial plane of NPC is huge and each IR monomer of NPC suffers from large defocus change, we chose patch CTF estimation to determine the CTF model of each micrograph, so that we can use linear interpolation to obtain the local CTF information of each IR monomer and IR dimer based on the CTF model.

**NPC particle picking**

We used a convolutional neural network (CNN)-based particle picking software, Topaz [53], which can accurately pick the particles including side-view particles and exclude the obvious bad particles. 1,731 manual-picked particles from 2,000 micrographs were used to train a particle picking model in Topaz. Because the diameter of the whole NPC particle is very large, in order to speed up the training speed, we downsample the micrographs by a downsampling factor of 105, which corresponds to the size of the convolution kernel. Finally, 279,900 particles were auto-picked from 187,076 micrographs.

**Generation of the initial model of NPC**

The initial model of NPC was generated in cryoSPARC. A subset of 123,769 particles from 279,900 particles were extracted with a box size of 512 pixels (binned 4x, 2.672 Å/pixel). After 2D classification, 11,624 particles were selected to generate an initial model. First, EMD-7321 was low-pass filtered to 30 Å and used as the initial model for Homogeneous Refinement. Then we subtracted the cargo signal from the particles according to the parameters obtained from the refinement. The subtracted particles were subjected to Ab-initio Reconstruction to get 7 models, followed by Heterogeneous Refinement applied with C8 symmetry. Finally, the model with the best resolution (22 Å) was chosen as the initial model.

**Whole NPC complex reconstruction**

The image processing was performed in cryoSPARC and RELION3.0/3.1 [54]. 279,900 particles were extracted with a box size of 512 pixels (binned 4x, 2.672 Å/pixel). After 2D classification in cryoSPARC, the obvious bad particles were removed. The remained 263,477 particles were imported into RELION3.0 for 3D classification and refinement. 3D classification was performed with global search, C8 symmetry, K=1 and 50 iterations, and the initial model generated above was low-pass filtered to 30 Å. Then another round of 3D classification with local search, C1 symmetry and K=1 was performed. By this step, we obtained one class showing eight spokes clearly even without applying symmetry. Then the particles from this class were submitted to 3D auto-refine with C8 symmetry to further improve the resolution. The resulted data star file was imported to cryoSPARC to re-extract particles followed by Non-uniform Refinement [55]. After this step, the signal of center cargo was subtracted from particles by particle subtraction. 2D classification of the subtracted particles indicated that the density of the center cargo was greatly reduced. Finally, we got a whole NPC map at 12.03 Å after Non-uniform Refinement with C8 symmetry imposed.

**IR reconstruction**

When we gradually increased the threshold of the whole NPC map, only IR region becomes clearly visible among the three layers, indicating that IR is the most stable and uniform region in the entire scNPC. So, we focused on the refinement of IR. Due to the huge diameter, the particle suffers from the large defocus change along the equatorial plane. In order to alleviate this problem, we tried a strategy: refine the CTF and pose parameters of one IR monomer for each NPC particle, and then use the refined parameters to reconstruct the whole IR without alignment. Firstly, local-refinement with an IR monomer mask was performed to further optimize the pose parameters. Then the particles were re-centered to the IR monomer using the refined pose parameters, and 278,938 particles were extracted with a 512-pixel box size (binned 2x, 1.336 Å/pixel) for the IR monomer. After 2D classification, the particles were divided into one good group and one bad group, which were used to generate one good and two bad initial models, respectively. Then, the total 278,938 particles were subjected to Heterogeneous Refinement with the three initial models generated above. The class with best quality was further subjected to Non-uniform Refinement, which resulted in a 8.22 Å map with 101,757 particles. A round of CTF refinement and Non-uniform Refinement was performed. Finally, the resolution was improved to 7.41 Å. Then, 101,757 particles were re-centered to the center of the whole NPC and extracted with a 512-pixel box size (binned 4x, 2.672 Å/pixel) for the whole NPC. The refined CTF parameters and pose information obtained from the 7.41 Å map of the IR monomer were applied back to the corresponding whole NPC particles for reconstruction without alignment and with an IR mask and C8 symmetry imposed, which resulted in a 13 Å IR map. After post-processing with an IR mask and B-factor of -800, we obtained an entire IR map at 9.10 Å.

**IR monomer/protomer and IR dimer reconstruction**

In order to further improve the resolution of the IR monomer, we local refined each IR monomer of whole NPC, then used the corresponding pose parameters to extract each IR monomer particle with a 350-pixel box size (binned 2x, 1.336 Å/pixel). The extracted 2,238,689 particles were subjected to 2D classification to remove obvious bad particles that were used to generate four bad initial models by Ab-initio Reconstruction. The remained 1,954,403 particles were used to reconstruct a good initial model by Reconstruction only. A round of Non-uniform Refinement and CTF refinement was performed. Then Heterogeneous Refinement with five initial models generated above was performed to remove bad particles. 795,543 particles allocated to the best class were subjected to Non-uniform Refinement. The resolution was improved to 3.98 Å without applying symmetry and 3.84 Å with C2 symmetry imposed. Using the 3.84 Å map combined four bad initial models, the second round of Heterogeneous Refinement was performed to screen out more bad particles from the initial 1,954,403 particles. 679,756 particles of the best class were selected for 2D classification and Non-uniform Refinement. We obtained a 3.73 Å map with 633,134 particles. In order to further improve the resolution to aid model building, we performed symmetry expansion of each IR monomer particle. Each monomer was replicated and rotated two times around its C2 (pseudo) symmetry axis, followed by signal subtraction to retain one copy of each protomer, and subsequent local refinement. Although the final resolution of protomer is 3.71 Å with twofold particles, only slightly higher than that of IR monomer, the map quality of CNT is much better than that in the IR monomer.

In order to improve the quality of the density at the connection region between adjacent IR monomers, we used the similar methods as above to obtain an IR dimer map at 7.69 Å.

All reported resolutions were estimated based on the gold-standard Fourier Shell Correlation (FSC) 0.143 criterion [54]. All local resolution maps were determined using cryoSPARC.

**Image processing of Nup157, Nup170 and Nup188**

For Nup157, a small dataset of 88,788 particles were picked from 200 micrographs using blob picker with particle diameter ranged from 70 Å to 200 Å and processed by reference-free 2D classification using cryoSPARC. 8,421 good particles with clear 2D averages were selected as training dataset for Topaz, by which a total of 2,652,917 particles were automatically picked from all micrographs. After 3 rounds of 2D classification, 1,993,084 particles were selected and subjected to 2 rounds of Ab-initio Reconstruction with 6 classes in cryoSPARC. Among these classes in the second round, class I with the clearest and the most complete density was selected for Homogeneous Refinement in cryoSPARC. Another round of 2D classification was executed for removing the incomplete particles lacking C-terminal region, and 100,523 particles were selected and imported into RELION3.0. After 3D auto-refinement and post-processing, the final resolution of the full length Nup157 density map is 5.9 Å. To analyze the particle heterogeneity, the 100,523 particles were imported into EMAN2.91 [56], and subjected to 2D classification. One class was chosen as the 2D reference. All the selected particles were aligned to the single 2D reference using e2a2d_align.py command, and further analyzed by e2motion.py command. The N-terminal region were used for realignment of the particles and the C terminal region were used for classification. Finally, 32 images of 2D class average were generated and used to make the animated movie (Supplementary Movie 3).

For Nup170, a small dataset of 59,240 particles were picked from 200 micrographs using blob picker with particle diameter ranged from 70 Å to 200 Å and processed by reference-free 2D classification using cryoSPARC. 6,700 good particles with clear 2D averages were selected as training dataset for Topaz, by which a total of 1,733,609 particles were automatically picked from all micrographs. After 2 rounds of 2D classification, 944,829 particles were selected and subjected to Ab-initio Reconstruction with 5 classes in cryoSPARC. We cannot get full length Nup170 map after trying many methods, and one class (130,369 particles) with the clearest density was selected for Non-uniform Refinement in cryoSPARC. Then particles were imported into RELION3.0, and CTF refinement, 3D auto-refinement and post-processing were applied. The final resolution of the Nup170 density map is 3.7 Å.

For Nup188, 12,050 micrographs were selected with the maximum resolution value below 4 Å and the astigmatism below 1000 Å. The following image processing steps were carried out in cryoSPARC and RELION 3.1 showed in Supplementary information, Fig. S12. 5,610,043 particles were auto-picked using blob picker with particle diameter ranged from 100 Å to 170 Å and subjected to 2D classification. 1,144,198 particles with good class-averages were selected and subjected to Ab-initio Reconstruction for 10 classes without reference. 2 classes (672,071 particles) with good structure features were selected and perform another Ab-initio Reconstruction for 8 classes without reference. 5 classes (607,216 particles) with good structure features were selected and subjected to Non-uniform Refinement with one good initial structure reconstructed above as reference, leading to a 3D map at an overall resolution of 2.81 Å. All these particles were imported into RELION 3.1, and auto-refined using the 2.81 Å map low-pass filtered to 60 Å as reference to reconstruct a map at an overall resolution of 3.02 Å after post-processing. A soft mask was applied for further auto-refine, and a 3D reconstruction at an overall resolution of 2.86 Å was obtained after post-processing. In the map, the density of the C-terminal part (amino acid 1216-1653) was much weaker than the N-terminal part.

To analyze the particle heterogeneity, all the 607,216 particles were imported into EMAN2.91 [56], and subjected to 2D classification. All top view classes (148,412 particles) with clear secondary structure features were selected and one of them was chosen as the 2D reference. All the selected particles were aligned to the single 2D reference using e2a2d_align.py command, and further analyzed by e2motion.py command. The N-terminal region were used for realignment of the particles and the C terminal region were used for classification. Finally, 48 images of 2D class average were generated and 46 images of them were selected to make the animated movie (Supplementary Movie 1).

To describe the continuous structural heterogeneity of C-terminal region in Nup188, two soft masks were made for N-terminal and C-terminal regions according to the 3.02 Å map low-pass filtered to 15 Å, respectively. All the 607,216 particles were subjected to multi-body refinement in RELION 3.1, and 3.02 Å and 3.41 Å reconstructions for the N-terminal and C-terminal regions were yielded, respectively (Supplementary information, Fig. S12). The movie of C-terminal region motions was generated (Movie 2).

All reported resolutions were estimated based on the gold-standard FSC 0.143 criterion. All local resolutions were calculated using RELION3.0.

**Model building and refinement**

Atomic coordinates of Nup188 were generated utilizing map-to-model software in PHENIX [57] with homologous crystal structures as the templates, including the N-terminus (PDB 5CWU and 4KF7) and C-terminus (PDB 4KF8), and manually refined in COOT [58] based on density of Nup188 determined in this study. The crystal structure of N-terminus (PDB 4MHC) and the predicted C-terminal model of Nup157 were combined and fitted into both the 5.9 Å map of the full-length Nup157 and the 3.73 Å map of the IR monomer to gain a composite model of Nup157. The N-terminal model generated by SWISS-MODEL [59] and the C-terminal crystal structure (PDB 3I5P) of Nup170 were manually adjusted based on the maps of Nup170, IR monomer and IR dimer to gain a composite model of Nup170. Model of Nic96 was from crystal structure (PDB 2RFO) lacking N-terminal disordered region. The atomic models of Nup192 and CNT complex were generated through homology modeling using SWISS-MODEL and I-TASSER [60] and adjusted based on the EM density map of IR monomer. All atomic coordinates for the individual subunits of IR were firstly fitted into the EM density map of the IR monomer based on the IR model from [18] (PDBDEV_00000051) using UCSF Chimera [61] and Chimera X [62] and then manually adjusted according to the EM density maps of IR monomer and IR dimer in COOT. Real space refinement and final validation of IR monomer model were performed through real space refine and validation program from PHENIX. The final atomic model of Nup188 was cross-validated according to previously described procedures [63]. Briefly, atoms in the model were randomly shifted by up to 0.5 Å, and then refined against one of the two independent half maps generated during the final 3D reconstruction. Then, the refined model was tested against the other map. FSC curves of the refined model versus the overall map (sum, blue), of the model refined against the first half map versus that same map (work, red), and of the model refined against the first half map versus the second map (free, green) (Supplementary information, Fig. S12). Structural visualizations and Figures were performed by UCSF Chimera, Chimera X and PyMOL (www.pymol.org).

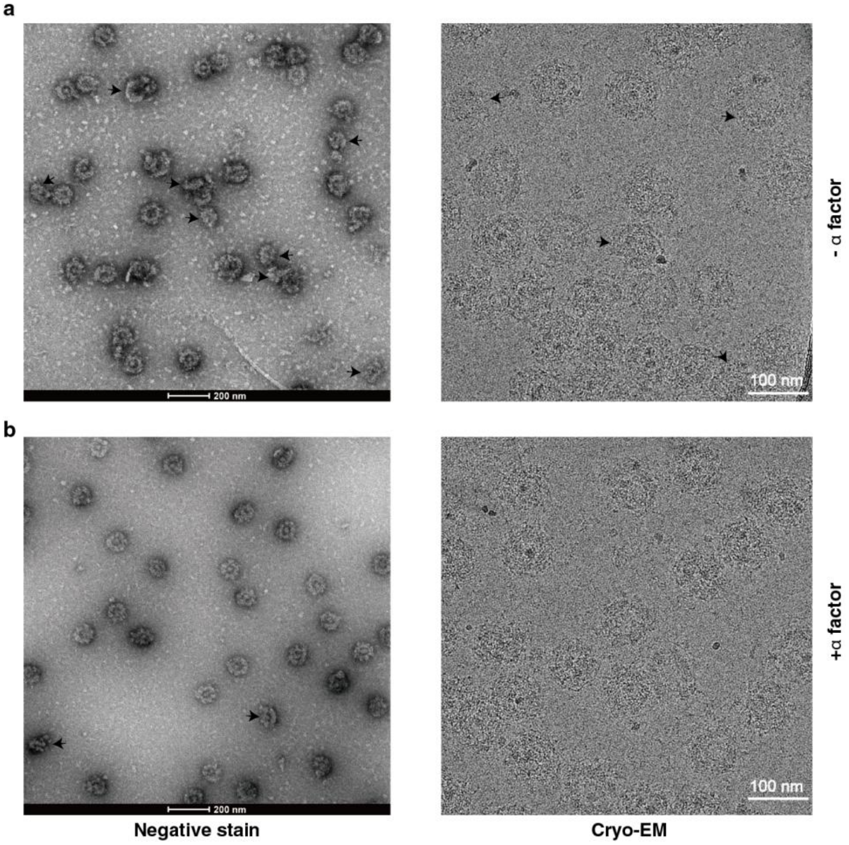


**Supplementary information, Fig. S1. Structural variability of *sc*NPC.**
(**a**, **b**) Representative negative staining EM and cryo-EM micrographs of *sc*NPC treated with or without alpha-mating factor. Heterogeneous particles are indicated by black arrows.

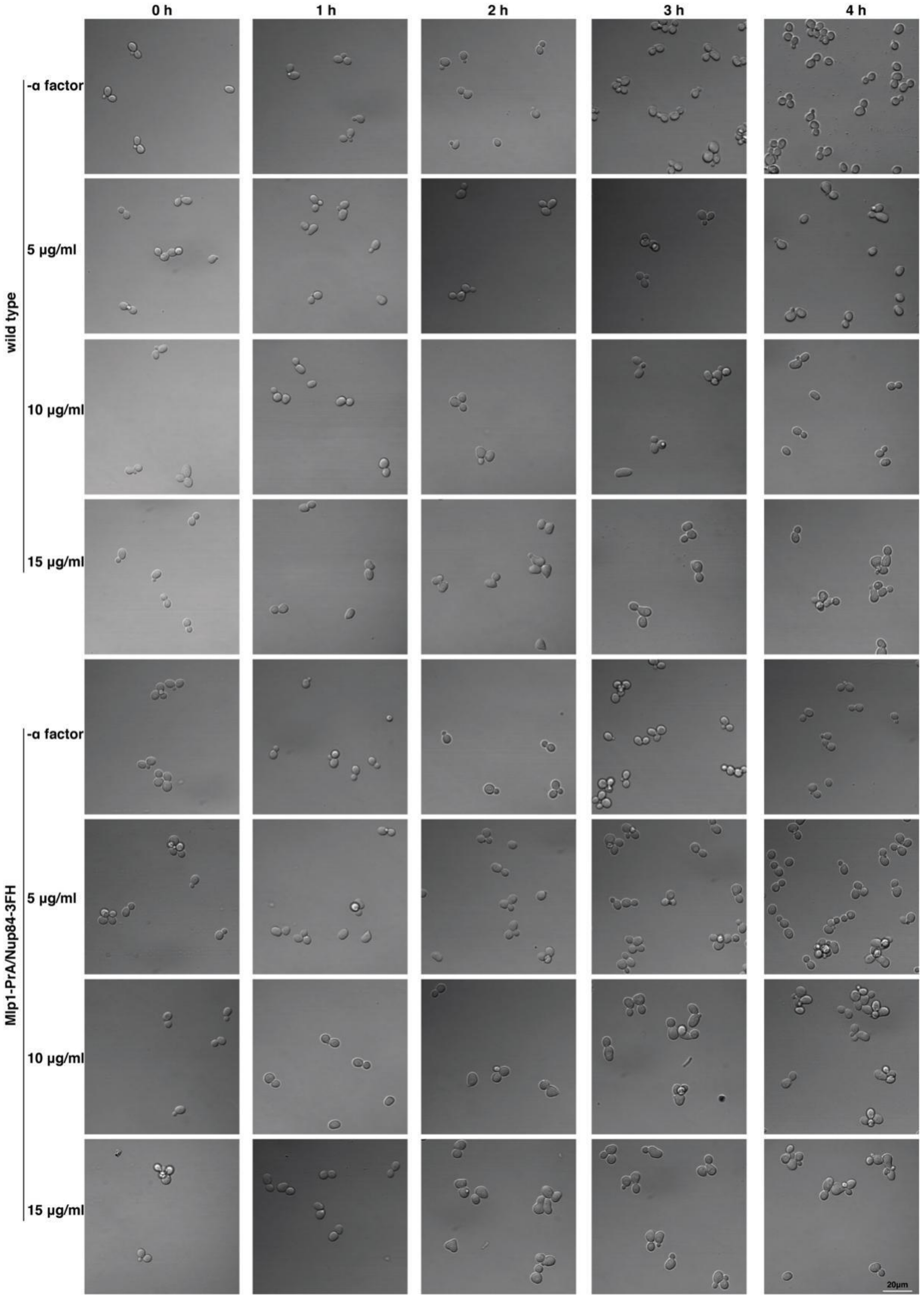


**Supplementary information, Fig. S2. Effect of alpha-mating factor on yeast cells.** Alpha-mating factor induced morphologic changes of wild-type and *Mlp1-PrA/Nup84-3FH* yeast cells. Different concentrations and treatment times were experimented and observed on laser scanning confocal microscope (Olympus FV1200). Shmoo shapes suggest cells are arrested in G1 phase.

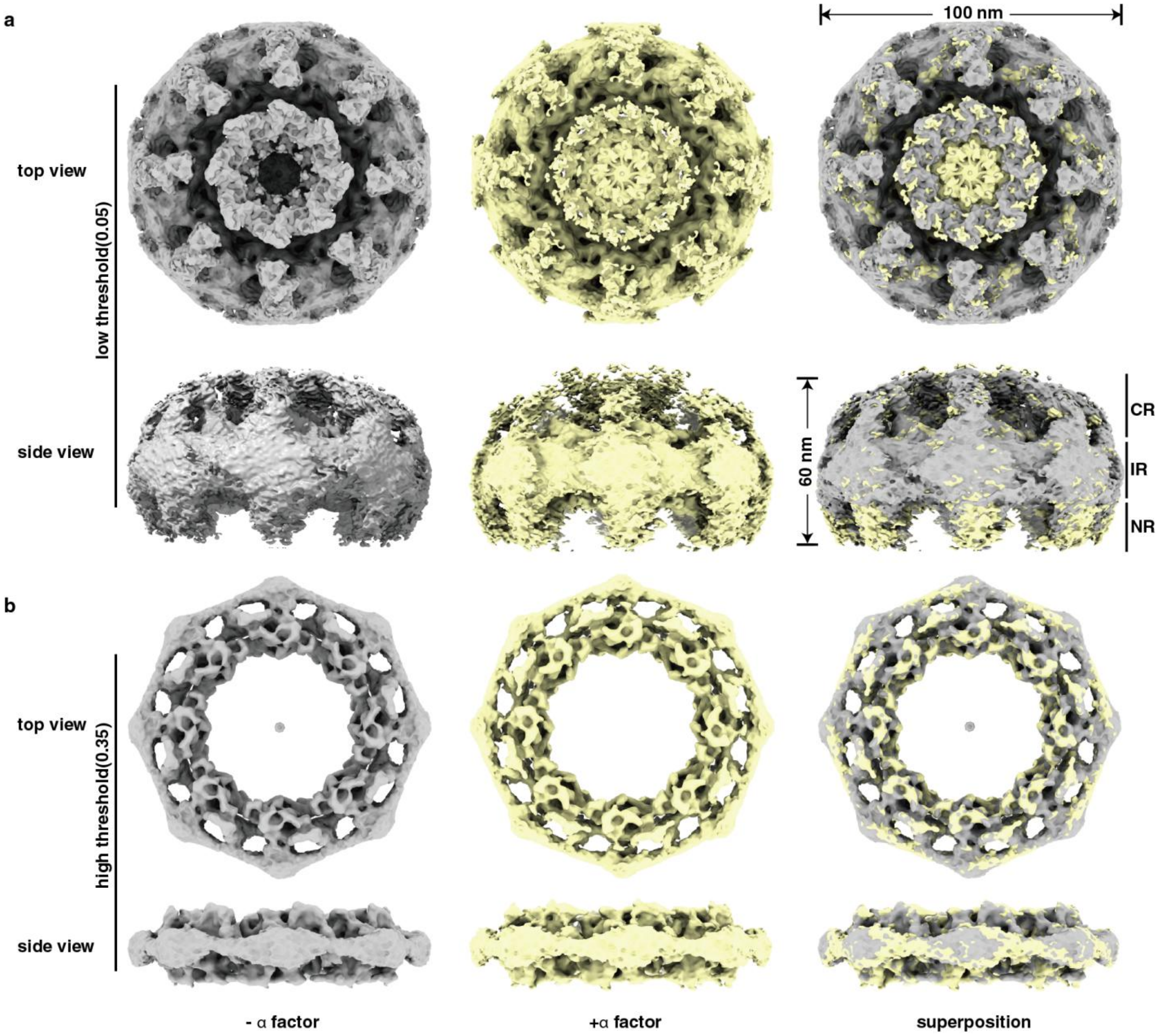


**Supplementary information, Fig. S3. Structures of NPC extracted from yeast cells treated with or without alpha-mating factor.**

(**a**, **b**) EM density maps are shown in low (a) and high (b) threshold, and superpositions are shown in the right panels.

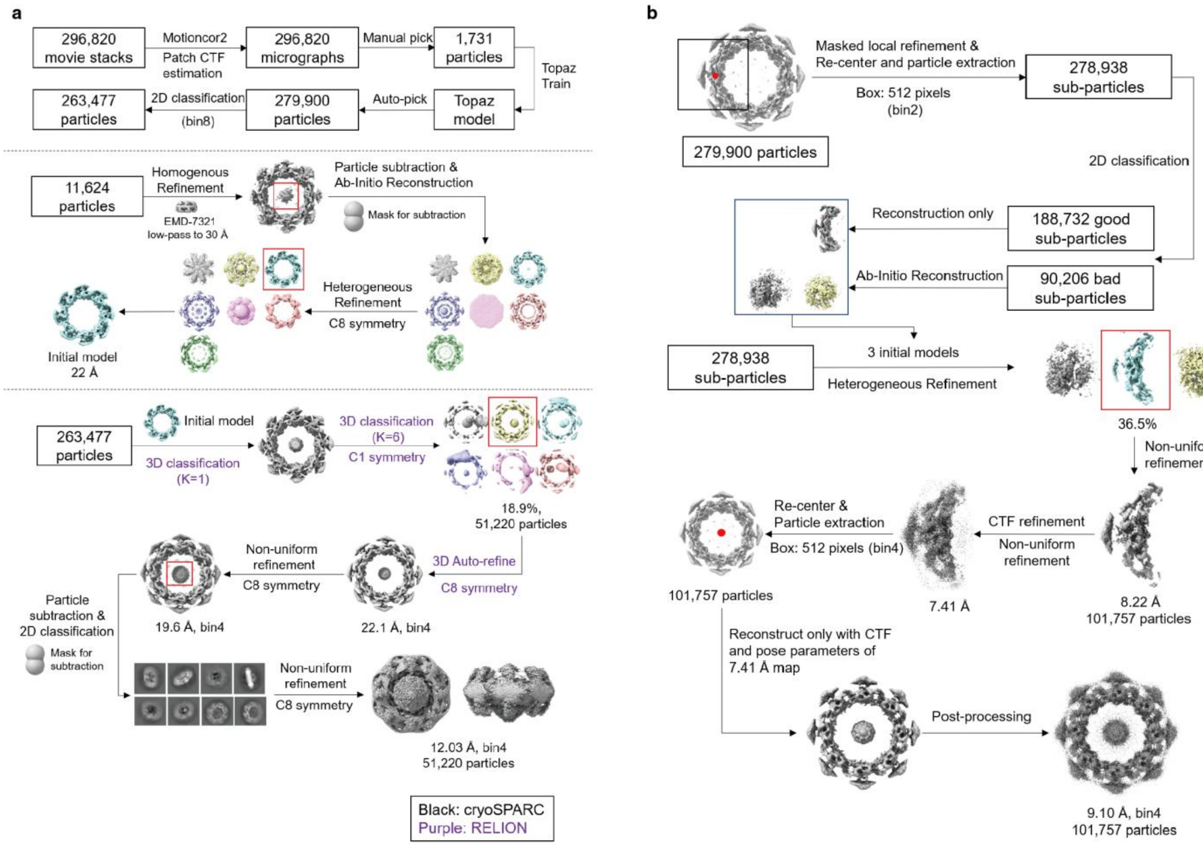


**Supplementary information, Fig. S4. Flowcharts for cryo-EM data processing of entire NPC (a) and entire IR (b).**
See “Materials and Methods” for details.

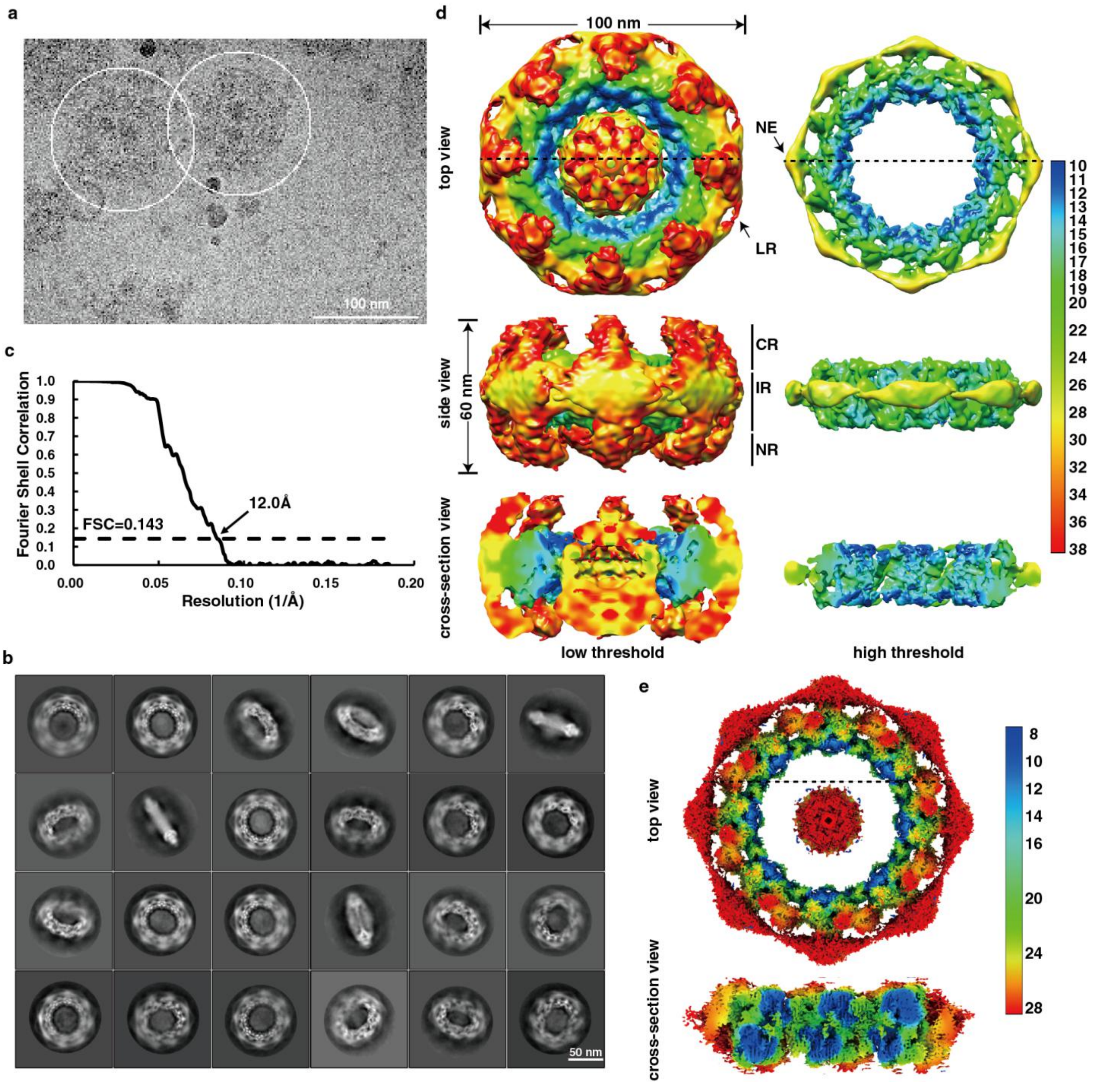


**Supplementary information, Fig. S5. Cryo-EM data analysis of entire NPC and entire IR.** (**a**) A representative raw cryo-EM image of NPC. (**b**) Typical good reference-free 2D class averages of NPC. (**c**) Gold standard FSC curve for the cryo-EM map of entire NPC. (**d**) Local resolutions of cryo-EM map for entire NPC with different threshold. (**e**) Local resolutions of cryo-EM map for intact IR at different views. Dotted line indicates the cross section corresponding to the side view.

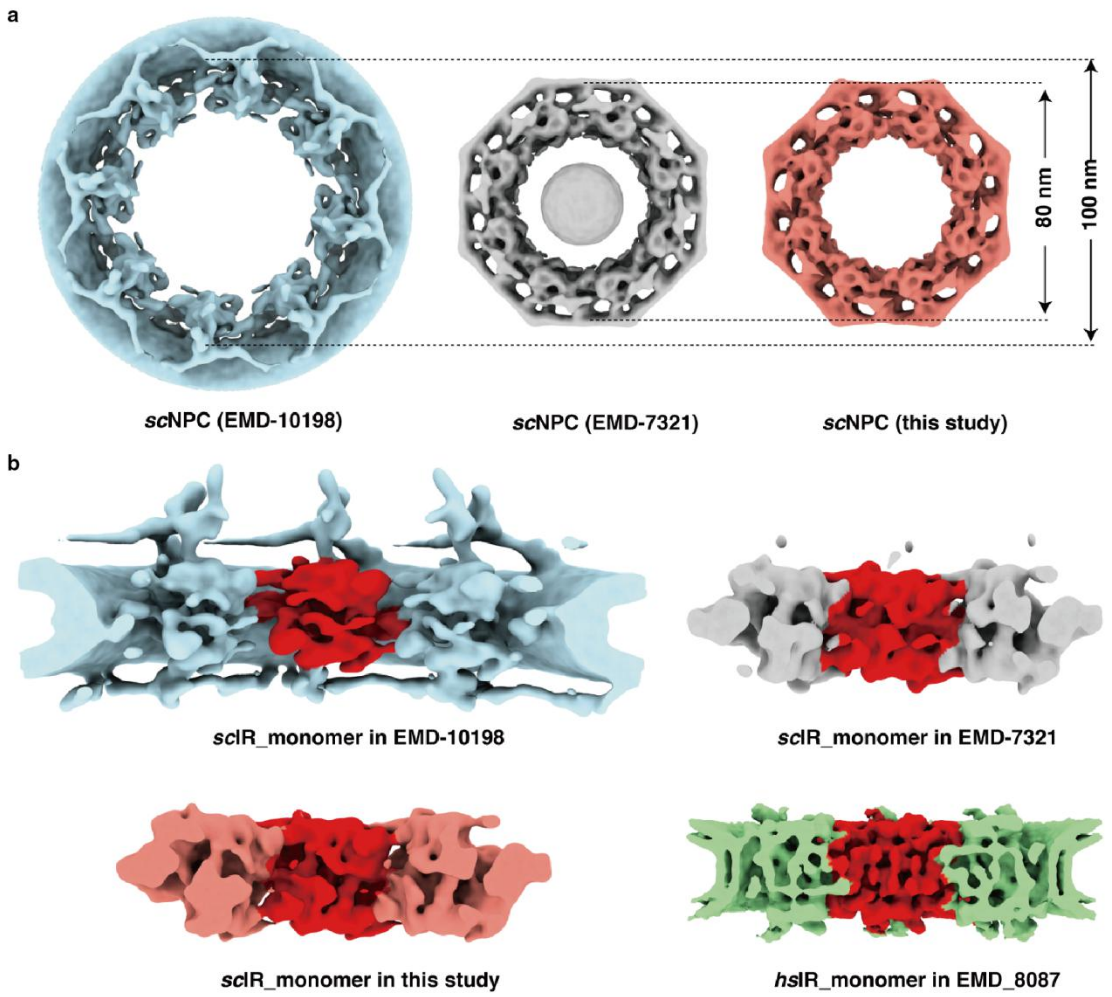


**Supplementary information, Fig. S6. The changeable intact IR and relatively stable IR monomer.**

(**a**) Comparison of *S. cerevisiae* IR from *in situ* architecture (EMD-10198) (34), detergent-extracted (EMD-7321) (27) and this study indicates ~ 20 nm diameter difference. (**b**) Comparison of IR monomer from different cell states and species including *S. cerevisiae* IR monomer at the active transport state (EMD-10198) and the static state (EMD-7321 and this study), and *Homo sapiens* IR monomer (EMD_8087).

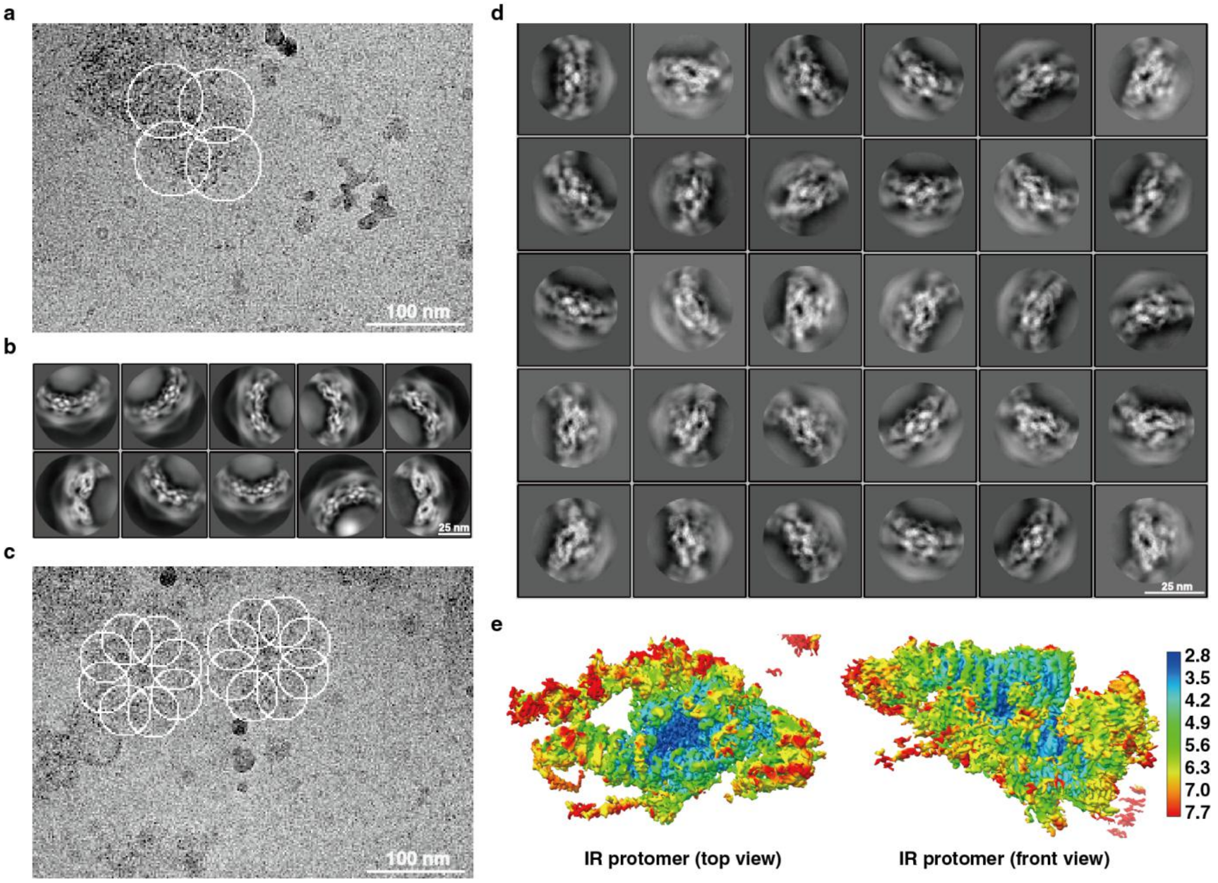


**Supplementary information, Fig. S7. Cryo-EM data analysis of IR dimer, monomer and protomer.**
(**a**) A representative raw cryo-EM image of NPC labeled with circles for IR dimer particle extraction. (**b**) Typical good reference-free 2D class averages of IR dimer. (**c**) A representative raw cryo-EM image of NPC labeled with circles for IR monomer particle extraction. (**d**) Typical good reference-free 2D class averages of IR monomer. (**e**) Local resolutions of cryo-EM map for IR protomer at different views.

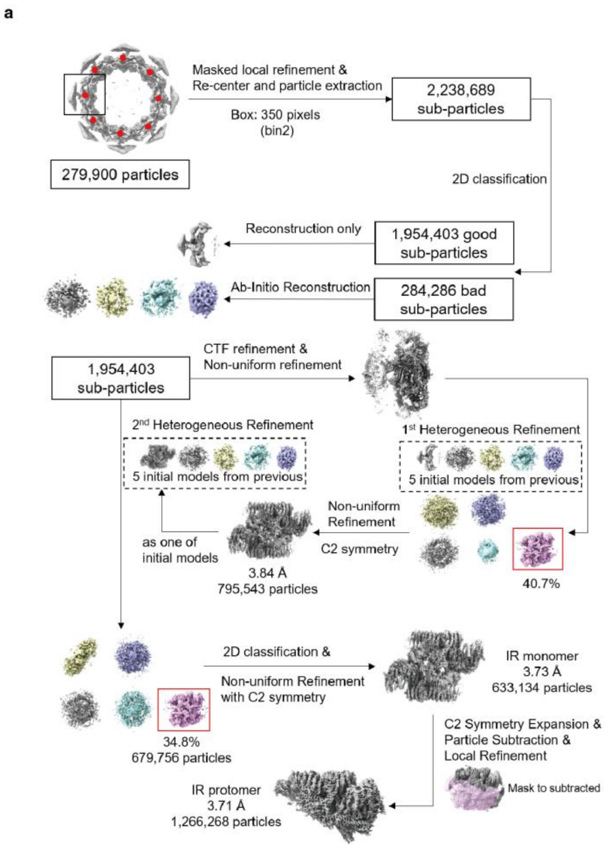


**Supplementary information, Fig. S8. Flowcharts for cryo-EM data processing of IR dimer (a), monomer and protomer (b)**.
See "Materials and Methods" for details.

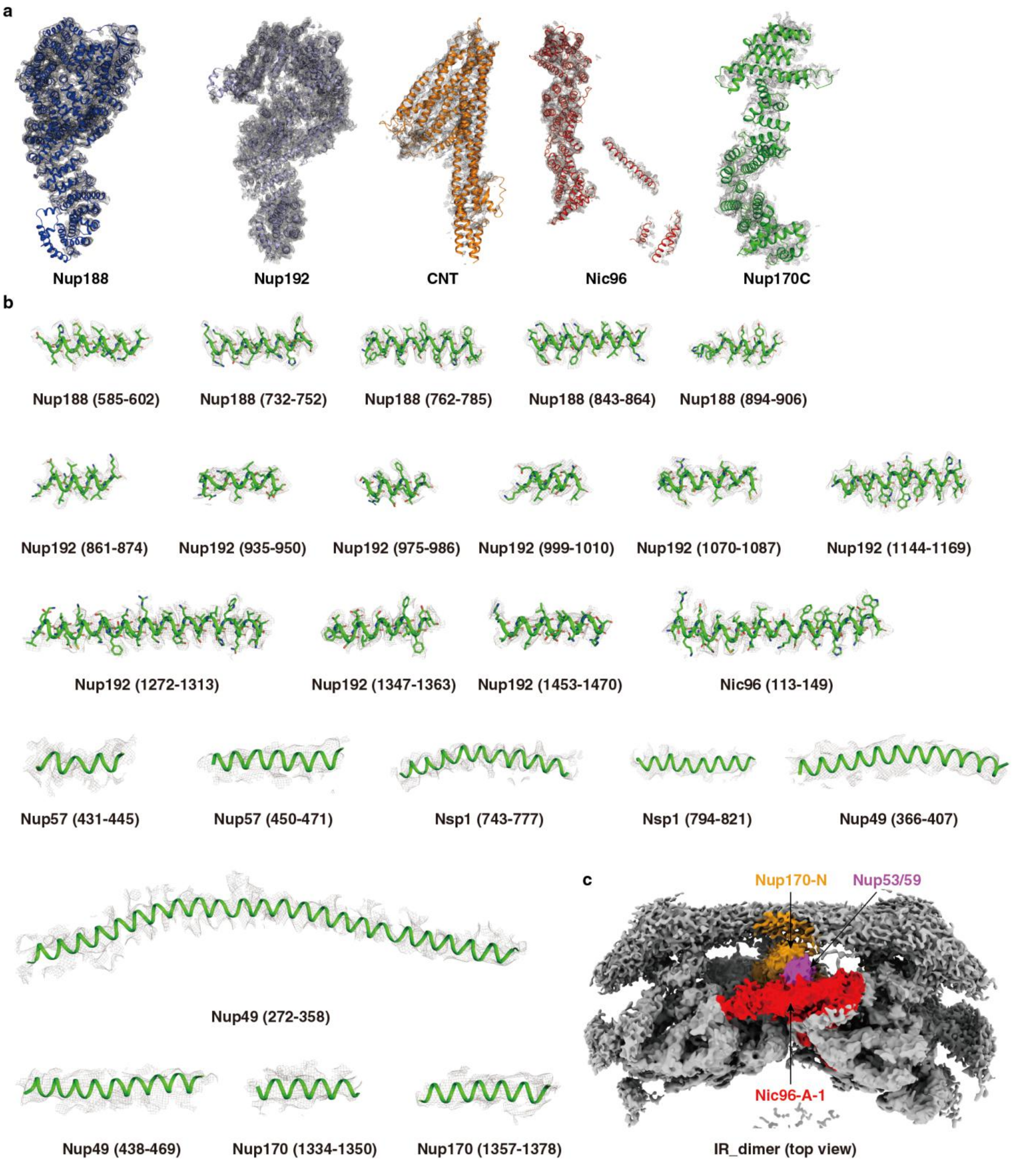


**Supplementary information, Fig. S9. Representative density maps of IR subunits.**
(**a**) The overall EM density map of IR subunits. (**b**) Representative EM density maps for a series of dis**c**rete α-helices from IR subunits. (**c**) Location of Nup53/Nup59 density based on our map and previously published biochemical results.

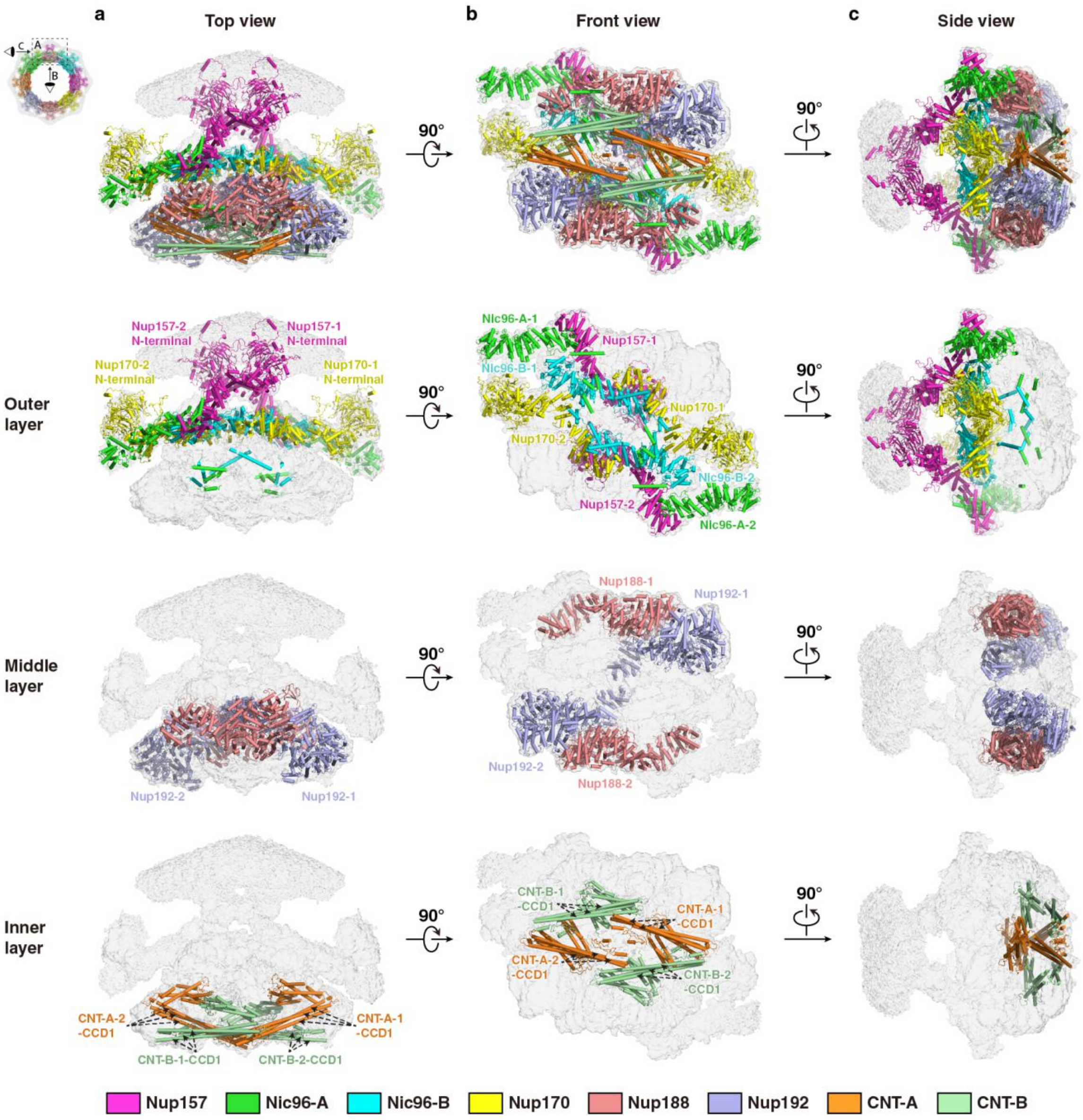


**Supplementary information, Fig. S10.Overall organization of the IR monomer.** (**a-c**) Overall structures of the IR monomer and each layer are shown in three perpendicular views. The IR monomer is divided into three layers: outer layer, middle layer and inner layer. The subunits of IR are color coded and shown in cartoon representation. IR monomer includes 24 proteins from 8 different Nups, Nup157, Nup170, Nic96, Nup88, Nup192, Nsp1, Nup49 and Nup57. The last three proteins (Nsp1, Nup49 and Nup57) form CNT complex.

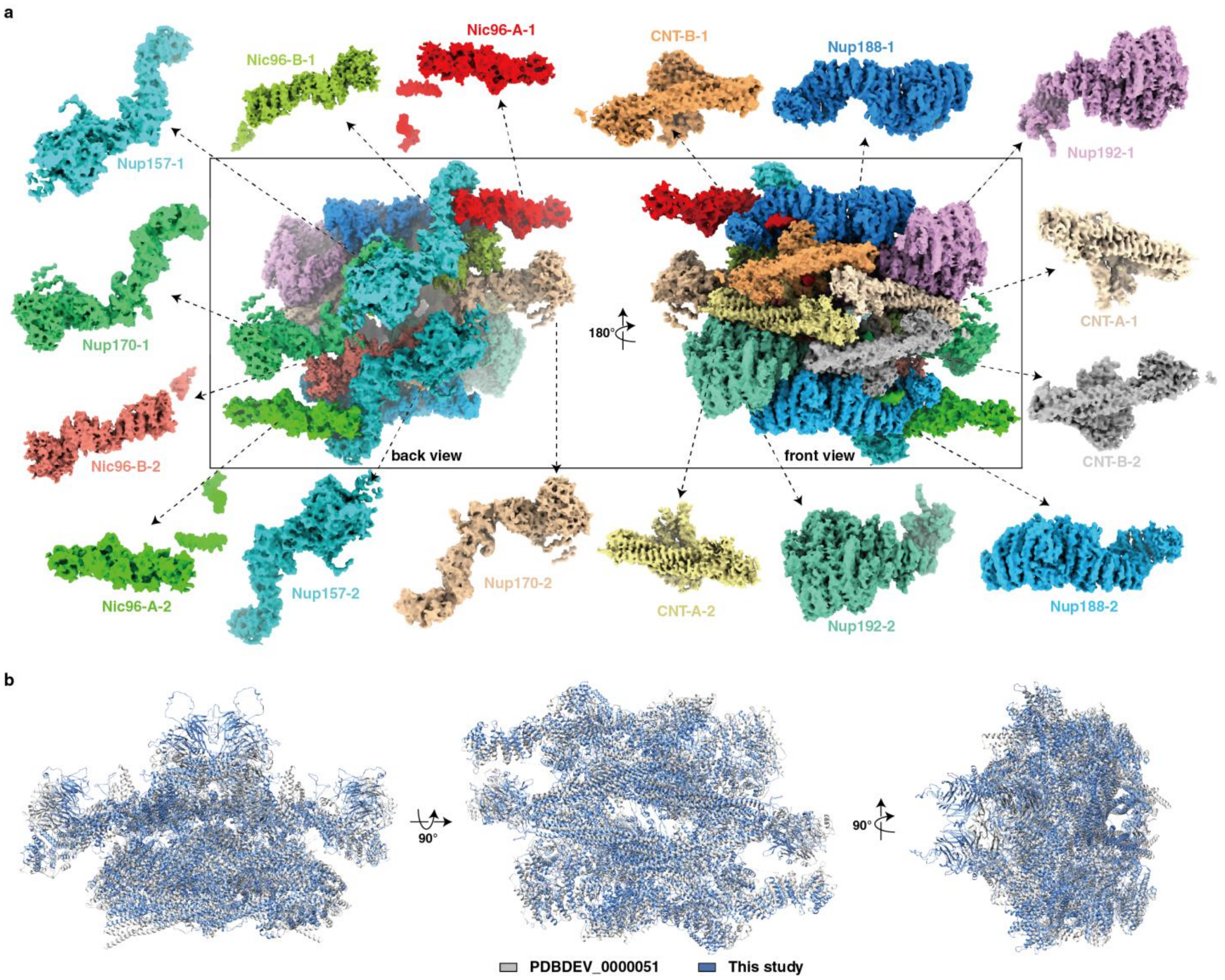


**Supplementary information, Fig. S11. Architecture of IR monomer.**
(**a**) The density map of IR monomer is shown in two views and 24 proteins from IR monomer are color-coded. CNT complex, which is composed of three proteins, Nup57, Nup49 and NSP1, is painted as the same color. (**b**) Structural comparison of IR monomer in this study and reported previously.

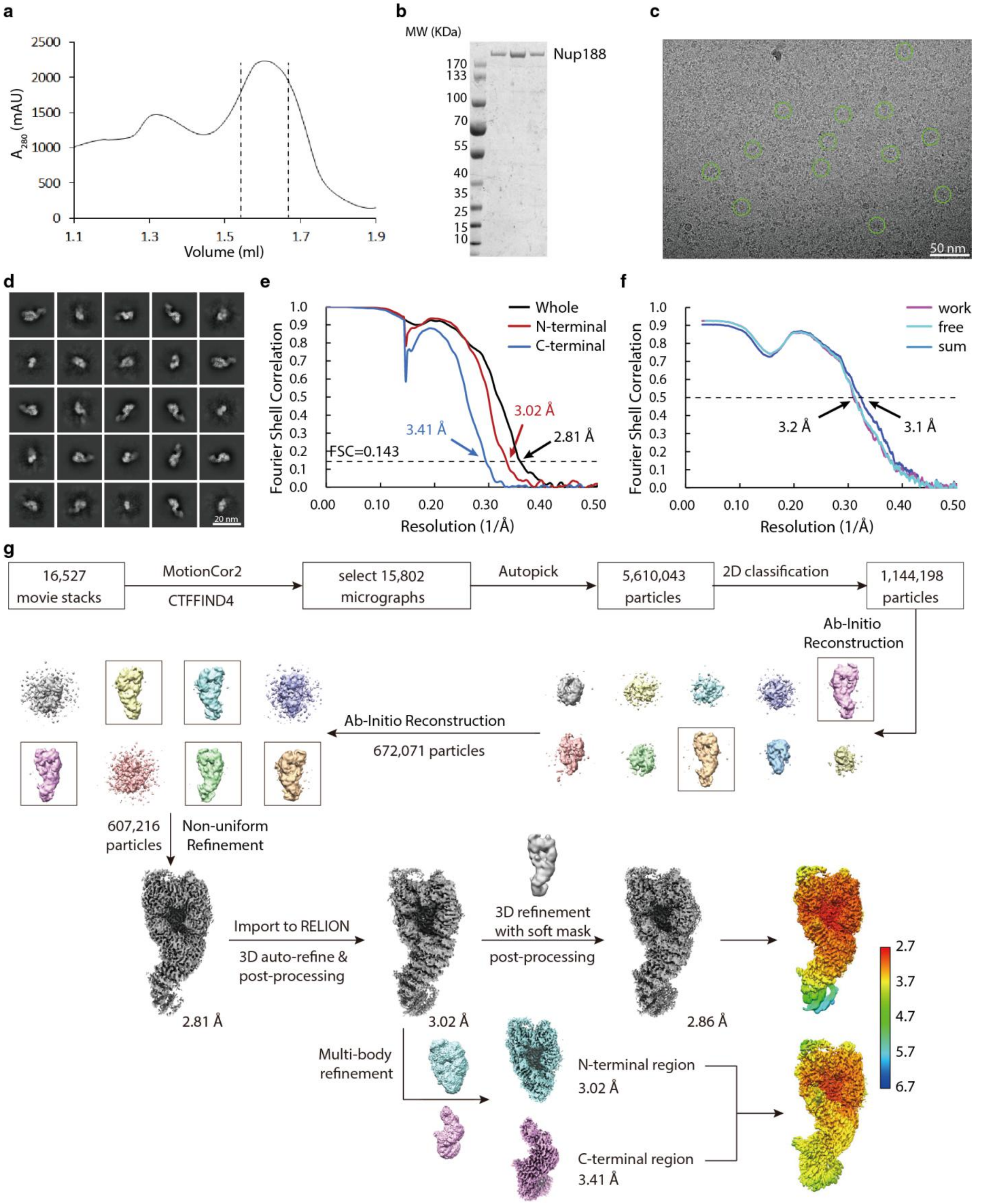


**Supplementary information, Fig. S12. Cryo-EM data analysis of Nup188.**
(**a**, **b**) Purification of Nup188. SEC profile of Nup188 (a) and the SDS-PAGE gel of the fractions corresponding to the region between dashed lines on SEC curve (b). (**c**) A representative raw

cryo-EM image for Nup188 with typical particles marked by green circles. (**d**) Typical good reference-free 2D class averages of Nup188. (**e**) Gold standard FSC curves for the cryo-EM maps of whole Nup188, its N-terminal part and its C-terminal part. (**f**) FSC curves of the cross-validation of the Nup188 model. See "Materials and Methods" for details. The small difference between the red and green curves indicates that the refinement of the atomic coordinates was not affected by overfitting. (**g**) The flowchart for EM data processing and the local-resolution maps. Details can be found in "Materials and Methods".

**a**

**Nup188**

**Nup188 1** MATPSFGNSSPQLTFTHVANFMNDAAADVSAVDAKQLAQIRQFLKANKTNLIESLNTIRQNVTSSGDHNKLRSTIANLLQINVDNDPFFA

**Nup188 91** QSEDLSHAVEFFMSERSSRLHIVYSLLVNPDIDLETYSFIDNDRFNVVGKLISIISSVIQNYDIITASSLAHDYNNDQDMFTIVSLVQLK

**Nup188 181** KFSDLKFILQILQILNLMILNTKVPVDIVNQWFLQYQNQFVEFCRNINSTDKSIDTSSLQLYKFQNFQDLSYLSETLISRISSLFTITTI

**Nup188 271** LILGLNTSIAQFDIQSPLYMDTETFDTVNSALENDVATNIVNEDPIFHPMIHYSWSFILYYRRALQSSESFDDSDITKFALFAESHDVLQ

**Nup188 361** KLNTLSEILSFDPVYTTVITVFLEFSLNFIPITASTSRVFAKIISKAPEQFIENFLTNDTFEKKLSIIKAKLPLLNESLIPLINLALIDT

**Nup188 451** EFANFELKDICSFAVTKSSLNDLDYDLIADTITNSSSSSDIIVPDLIELKSDLLVAPPLENENSNCLLSIPKSTKGKILTIKQQQQQQQQ

**Nup188 541** QNGQQPPTTSNLIIFLYKFNGWSLVGRILQNLLHSYMEKGTQLDDLQHELMISIIKLVTNVVDPKTSIEKSSEILSYLSNSLDTSASTIN

**Nup188 631** GASIIQVIFEIFEISLQRKDYTSIVQCCEFMTMLTPNYLHLVSSYLNKSDLLDKYGKTGLSNMILGSVELSTGDYTFTIQLLKLTKVFIR

**Nup188 721** ESLSLKNIHISKRSKIDIINKLILHAIHIFESYYNWKYNNFLQKFEIAFHLTLIFYDVLHDVFTINPHQKDQLIISSSANKLLQLFLTPM

**Nup188 811** DSIDLAPNTLTNILISPLNTTTKILGDKILGNLYSKVMNNSFKLCTLLIAIRGSNRDLKPSNLEKLLFINSSKLVDVYTLPSYVHFKVQI

**Nup188 901** ELLSYLVEAPWNDDYPFLLSFLGEAKSMAFLKEVLSDLSSPVQDWNLLRSLYIFFTTLLESKQDGLSILFLTGQFASNKKINDESSIDK

**Nup188 991** KSSILTVLQKNSLLLDSTPEEVSCKLLETITYVLNTWTNSKIFIKDPKFVNSLLAKLKDSKKLFQKKENLTRDETVSLIKKYKLISRIVE

**Nup188 1081** FALCIYNSTDSNSEILNFLNQEDLFELVHHFFQIDGFNKTFHDELNLKFKEKWPSLELQSFQKIPLSRINENENFGYDIPLLDIVLKAD

**Nup188 1171** RSWNEPSKSQTNFKEEITDASLNLQYVNYEISTAKAWGALITTFVKRSTVPLNDGFVDLVEHFLKLNIDFGSDKQMFTQIYLERIELSFY

**Nup188 1261** LYSFKLSGKLLKEEKIIELMNKIFTIFKSGEIDFIKNIGKSLKNNFYRPLLRSVLVLLELVSSGDRFIELISDQLLEFFELVFSKGVYL

**Nup188 1351** LSEILCQINKCSTRGLSTDHTTQIVNLEDNTQDLLLLLSLFKKITNVNPSKNFNVILASSLNEVGTLKVILNLYSSAHLIRINDEPILG

**Nup188 1441** QITLTFISELCSIEPIAAKLINSGLYSVLLESPLSVAIQQGDIKPEFSPRLHNIWSNGLLSIVLLLLSQFGIKVLPETCLFVSYFGKQIK

**Nup188 1531** STIYNWGDNKLAVSSSLIKETNQLVLLQKMLNLLNYQELFIQPKNSDDQQEAVELVIGLDSEHDKKRLSAALSKFLTHPKYLNSRIIPTT

**Nup188 1621** LEEQQQLEDESSRLEFVKGISRDIKALQDSLFKDV

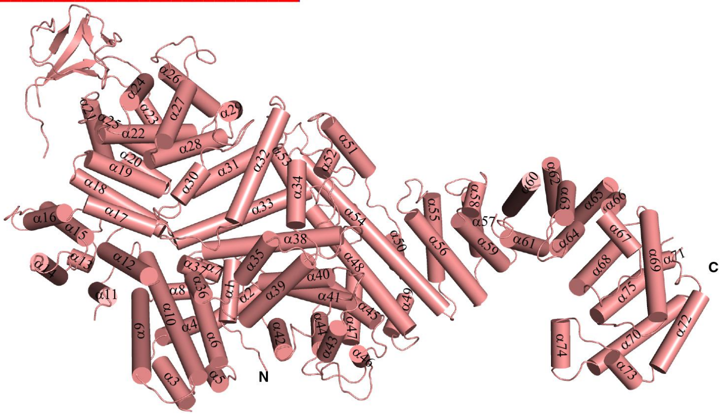

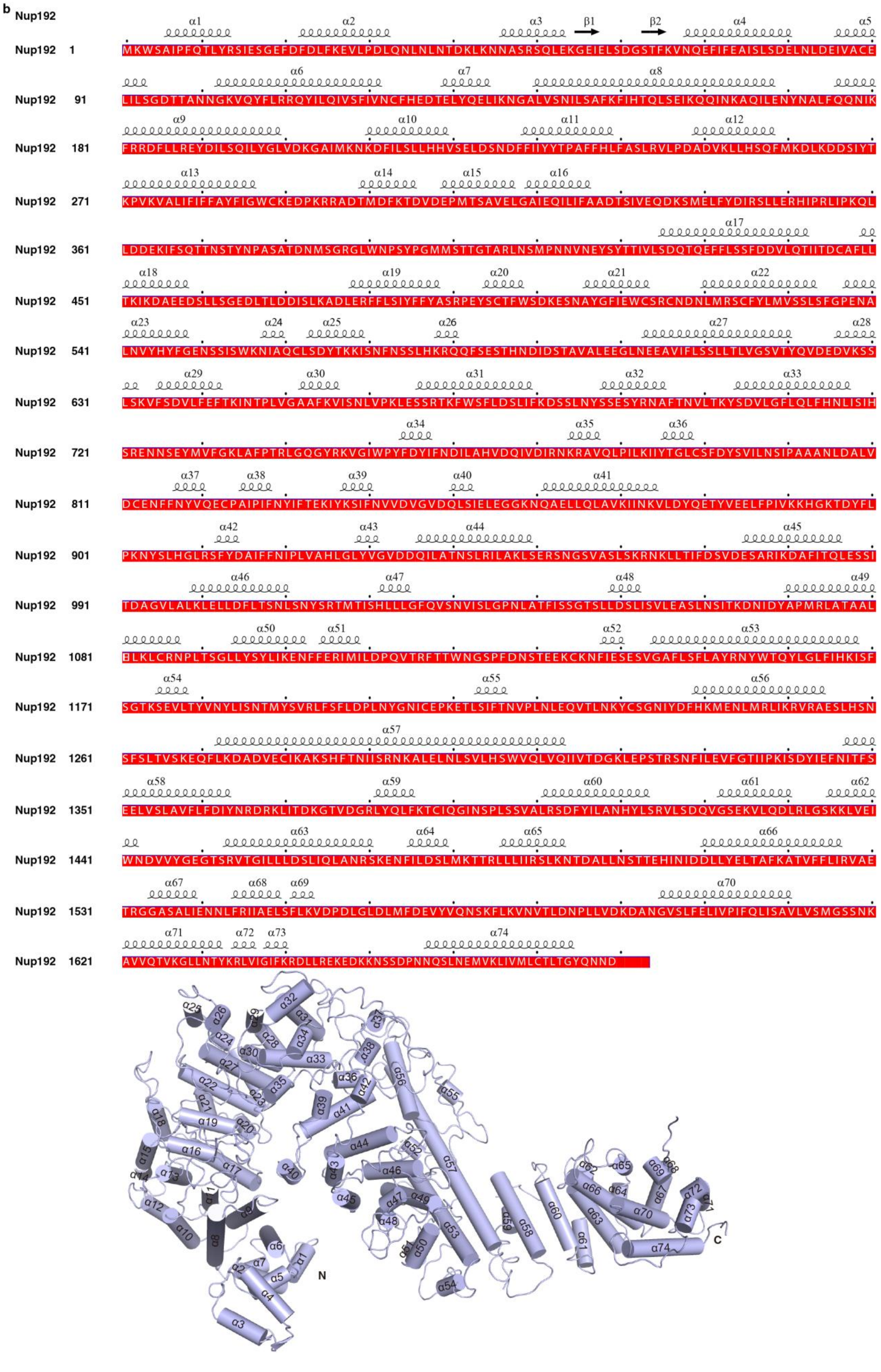
b
Nup192
Nup192 1
Nup192 91
Nup192 181
Nup192 271
Nup192 361
Nup192 451
Nup192 541
Nup192 631
Nup192 721
Nup192 811
Nup192 901
Nup192 991
Nup192 1081
Nup192 1171
Nup192 1261
Nup192 1351
Nup192 1441
Nup192 1531
Nup192 1621
N
C

c **Nup157**

α1

**Nup157 1** MYSTPLKKRIDYDRETFTASASLGGNRLRNRPRDDQNNGKPNLSSRSFLSERKTRKDVLNKYGEAGNTIESELRDVTTHVKISGLTSSEP

α2 α3 β1 β2 α4 β3 α5 β4

**Nup157 91** LQLASEFVQDLSFRDRNTPILDNPDYYSKGLDYNFSDEVGGLGAFTPFQRQQVTNIPDEVLSQVSNTEIKSDMGIFLELNYCWITSDNKL

β5 β6 β7 β8 β9 β10 β11

**Nup157 181** ILWNINNSSEYHCIDEIEHTILKVKLVKPSPNTFVSSVENLLIVATLFDIYILTISFNDRTHELNIFNTGLKVNVTGFNVSNIISYERTG

β12 β13 β14 α6 β15 β16

**Nup157 271** QIFFTGATDGVNVWELQYNCSENLFNSKSNKICLTKSNLANLLPTKLIPSIPGGKLIQKVLEGDAGTEEETISQLEVDQSRGVLHTLSTK

β17 β18 α7 β19 β20 β21 β22 β23 β24

**Nup157 361** SIVRSYLITSNGLVGPVLIDAAHIRRGMNALGVKNSPLLSNRAFKIAKIVSISMCENNDLFLAVITTTGVRLYFKGSISRRSIGSLKLDS

α8 α9 β25 β26 β27

**Nup157 451** VKFPPTSISSSLEQNKSFIIGHHPLNTHDTGPLSTQKASSTYINTTCASTIISPGIYFTCVRKRANSGELSKGITNKALLENKEEHKLYV

α10 β28 β29 α11 β30 β31 α12

**Nup157 541** SAPDYGILKNYGKYVENTALLDTTDEIKEIVPLTRSFNYTSTPQGYANVFASQYSAEPLKVAVLTSNALEIYCYRTPDEVFESLIENPLP

α13 α14 α15 α16 β32 α17

**Nup157 631** FIHSYGLSEACSTALYLACKFNKSEHIKSSALAFFSAGIPGVVEIKPKSSRESGSVPPISQNLFDKSGECDGIVLSPRFYGSALLITRLF

β34 β35 α18 α19

**Nup157 721** SQIWEERVFVFKRASKTEKMDAFGISITRPQVEYYLSSISVLADFFNIHRPSFVSFVPPKGSNAITASDAESIAMNALILLINSIKDALS

α20 α21 α22 α23 α24

**Nup157 811** LINVFYEDIDAFKSLLNTLMGAGGVYDSKTREYFFDLKFHDLFTPNAKTKQLIKEILIEVVNANIASGTSADYIVNVLKERFGSFCHSAD

α25 α26 α27 α28 α29

**Nup157 901** LCYRAGEHLEAAQKFEMIDSKISRNHLDTAIDLYERCAENIELCELRRVVDIMVKLNYQPKTVGFLLRFADKIDKGNQAQEYVSRGCNT

α30 α31 α32 α33

**Nup157 991** ADPRKVFYDKRINVYTLIFEIVKSVDDYTSIEQSPSIANISIFSPASSLKKRVYSVIMNSNNRFFHYCFYDWLVANKRQDYLLRLDSQFV

α34 α35 α36 α37 α38

**Nup157 1081** LPYLKERAEKSLEISNLLWFYLFKEEHFLEAADVLYALASSDFDLKLSERIECLARANGLCDSSTSFDQKPALVQLSENIHELFDIASIQ

α39 α40 α41 α42

**Nup157 1171** DDLLNLVRNETRIDEDYRKQLTLKLNGRVLPLSDLFNDCADPLDYYEIKLRIFKVSQFKDEKVIQGEWNRLLDSMKNAPSPDVGSVGQES

α43 α44 α45 α46 α47

**Nup157 1261** FLSSISNTLIRIGKTTRDTDVVFPVHFLMNKILESFIDKSSAADGSVCSMFLLAGVSHLKLYYILSRIIENSEGNVELAKKEMVWLIKDW

α48 α49 α50

**Nup157 1351** YQSDSDLRGSIAPEQIKKLEKYDPNTDPVQDYVKDRHHGLK

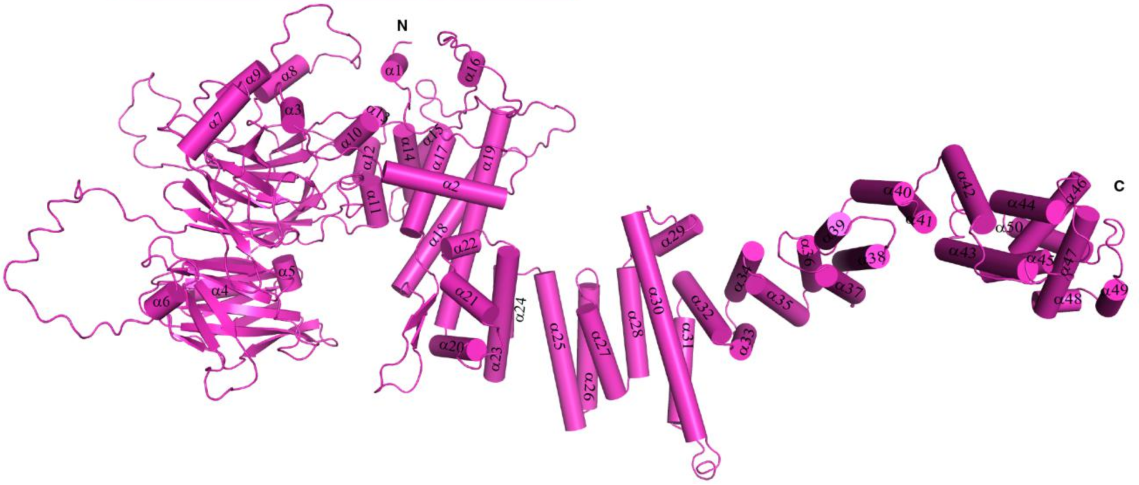

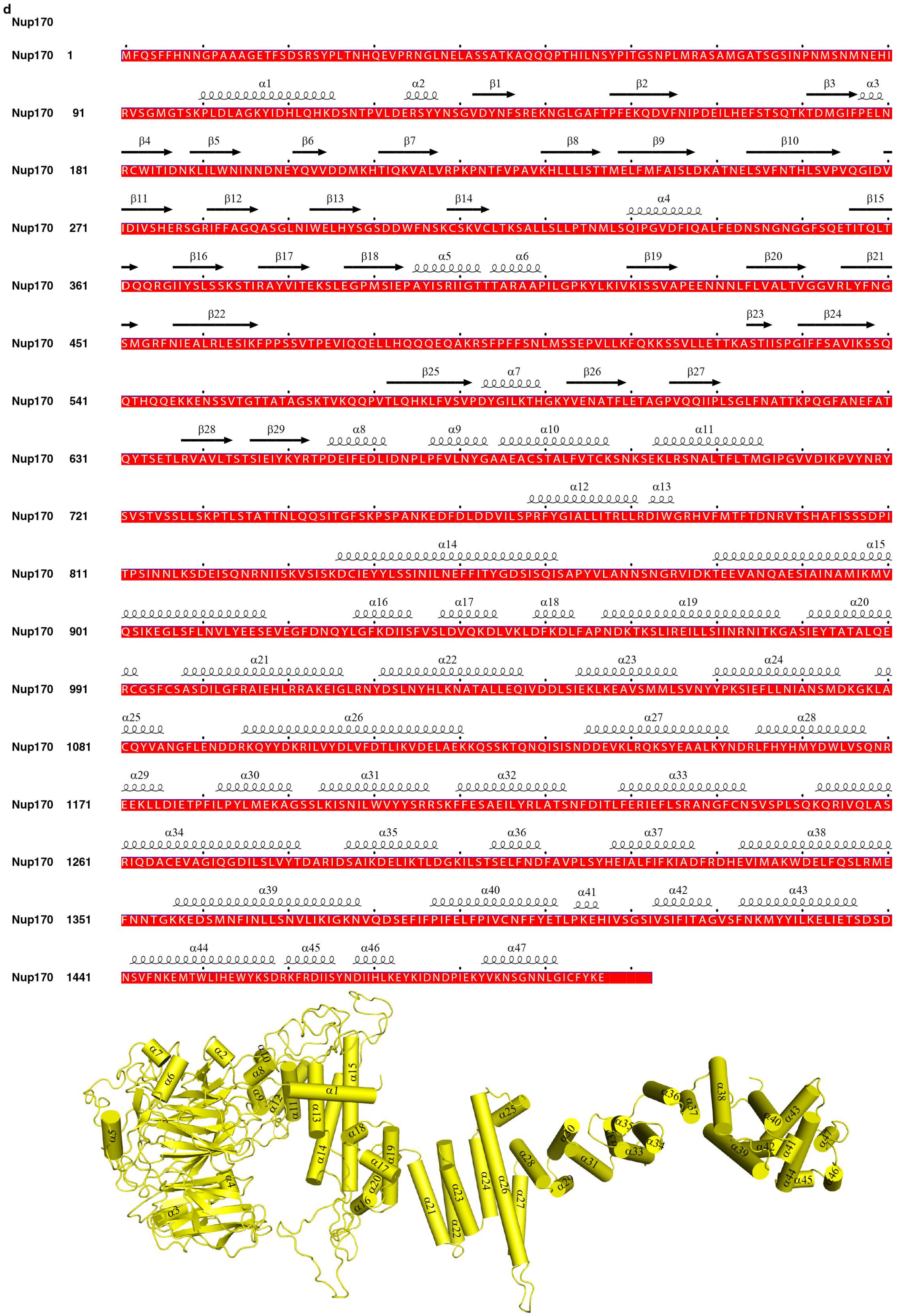
d
Nup170

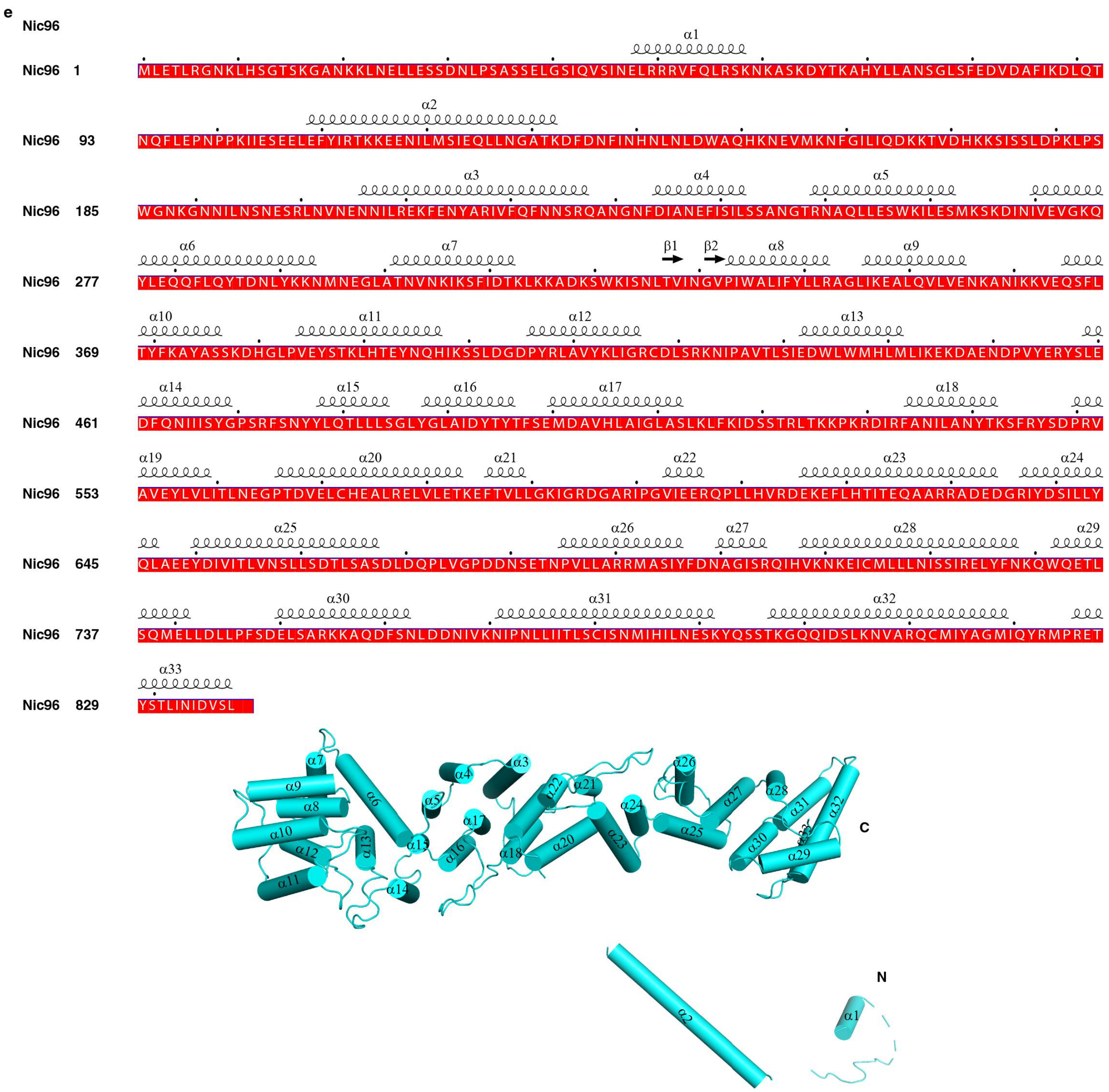
e
Nic96
α1
Nic96 1 MLETLRGNKLHSGTSKGANKKLNELLESSDNLPSASSELGSIQVSINELRRRVFQLRSKNKASKDYTKAHYLLANSGLSFEDVDAFIKDLQT
α2
Nic96 93 NQFLEPNPPKIIESEELEFYIRTKKEENILMSIEQLLNGATKDFDNFINHNLNLDWAQHKNEVMKNFGILIQDKKTVDHKKSISSLDPKLPS
α3 α4 α5
Nic96 185 WGNKGNNILNSNESRLNVNENNILREKFENYARIVFQFNNSRQANGNFDIANEFISILSSANGTRNAQLLESWKILESMKSKDINIVEVGKQ
α6 α7 β1 β2 α8 α9
Nic96 277 YLEQQFLQYTDNLYKKNMNEGLATNVNKIKSFIDTKLKKADKSWKISNLTVINGVPIWALIFYLLRAGLIKEALQVLVENKANIKKVEQSFL
α10 α11 α12 α13
Nic96 369 TYFKAYASSKDHGLPVEYSTKLHTEYNQHIKSSLDGDPYRLAVYKLIGRCDLSRKNIPAVTLSIEDWLWMHLMLIKEKDAENDPVYERYSLE
α14 α15 α16 α17 α18
Nic96 461 DFQNIIISYGPSRFSNYYLQTLLLSGLYGLAIDYTYTFSEMDAVHLAIGLASLKLFKIDSSTRLTKKPKRDIRFANILANYTKSFRYSDPRV
α19 α20 α21 α22 α23 α24
Nic96 553 AVEYLVLITLNEGPTDVELCHEALRELVLETKEFTVLLGKIGRDGARIPGVIEERQPLLHVRDEKEFLHTITEQAARRADEDGRIYDSILLY
α25 α26 α27 α28 α29
Nic96 645 QLAEEYDIVITLVNSLLSDTLSASDLDQPLVGPDDNSETNPVLLARRMASIYFDNAGISRQIHVKNKEICMLLLNISSIRELYFNKQWQETL
α30 α31 α32
Nic96 737 SQMELLDLLPFSDELSARKKAQDFSNLDDNIVKNIPNLLIITLSCISNMIHILNESKYQSSTKGQQIDSLKNVARQCMIYAGMIQYRMPRET
α33
Nic96 829 YSTLINIDVSL
C
N

**f**
**Nup57**

**Nup57 1** MFGFSGSNNGFGNKPAGSTGFSFGQNNNNTNTQPSASGFGFGGSQPNSGTATTGGFGANQATNTFGSNQQSSTGGGLFGNKPALGSLGSSST

**Nup57 93** TASGTTATGTGLFGQQTAQPQQSTIGGGLFGNKPTTTTGGLFGNSAQNNSTTSGGLFGNKVGSTGSLMGGNSTQNTSNMNAGGLFGAKPQNT

**Nup57 185** TATTGGLFGSKPQGSTTNGGLFGSGTQNNNTLGGGGLFGQSQQPQTNTAPGLGNTVSTQPSFAWSKPSTGSNLQQQQQQQIQVPLQQTAQAI

α1 α2

**Nup57 277** QQQQLSNYPQQIQEQVLKCKESWDPNTTKTKLRAFVYNKVNETEAILYTKPGHVLQEEWDQAMEKKPSPQTIPIQIYGFEGLNQRNQVQTEN

α3 α4

**Nup57 369** VAQARIILNHILEKSTQLQQKHELDTASRILKAQSRNVEIEKRILKLGTQLATLKNRGLPLGIAEEKMWSQFQTLLQRSEDPAGLGKTNELW

α5 α6 α7

**Nup57 461** ARLAILKERAKNISSQLDSKLMVFNDDTKNQDSMSKGTGEESNDRINKIVEILTNQQRGITYLNEVLEKDAAIVKKYKNKT

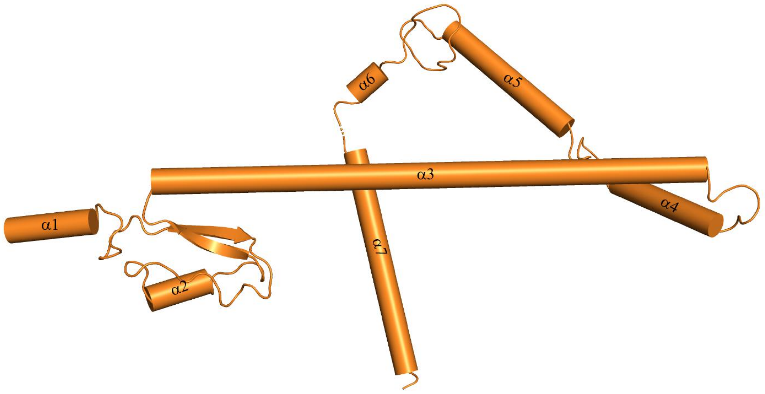


**g**
**Nup49**

**Nup49 1** MFGLNKASSTPAGGLFGQASGASTGNANTGFSFGGTQTGQNTGPSTGGLFGAKPAGSTGGLGASFGQQQQQSQTNAFGGSATTGGGLFGNKP

**Nup49 93** NNTANTGGGLFGANSNSNSGSLFGSNNAQTSRGLFGNNNTNNINNSSSGMNNASAGLFGSKPAGGTSLFGNTSTSSAPAQNQGMFGAKPAGT

**Nup49 185** SLFGNNAGNTTTGGGLFGSKPTGATSLFGSSNNNNNNNNSNNIMSASGGLFGNQQQQLQQQPQMQCALQNLSQLPITPMTRISELPPQIRQE

α1

**Nup49 277** EQLDQYIQKQVQISHHLKADTIDHDELIDSIPRDVAYLLKSESATSQYLKQDLKKISSFKSLIDEDLLDTQTFSVLLQQLLTPGSKISSND

α2 α3

**Nup49 369** LDKFFQKKIHLYEKKLEDYCRILSDIETAVNGIDTDLFGAPNNPNSTAITADLGSSEAENLLQLKTGLAAIVSTVIEEFTLFMDIAERIAVL

**Nup49 461** HQKTKTLASLSI

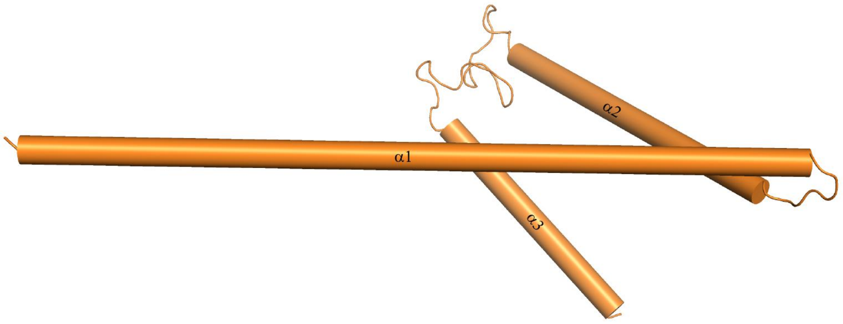

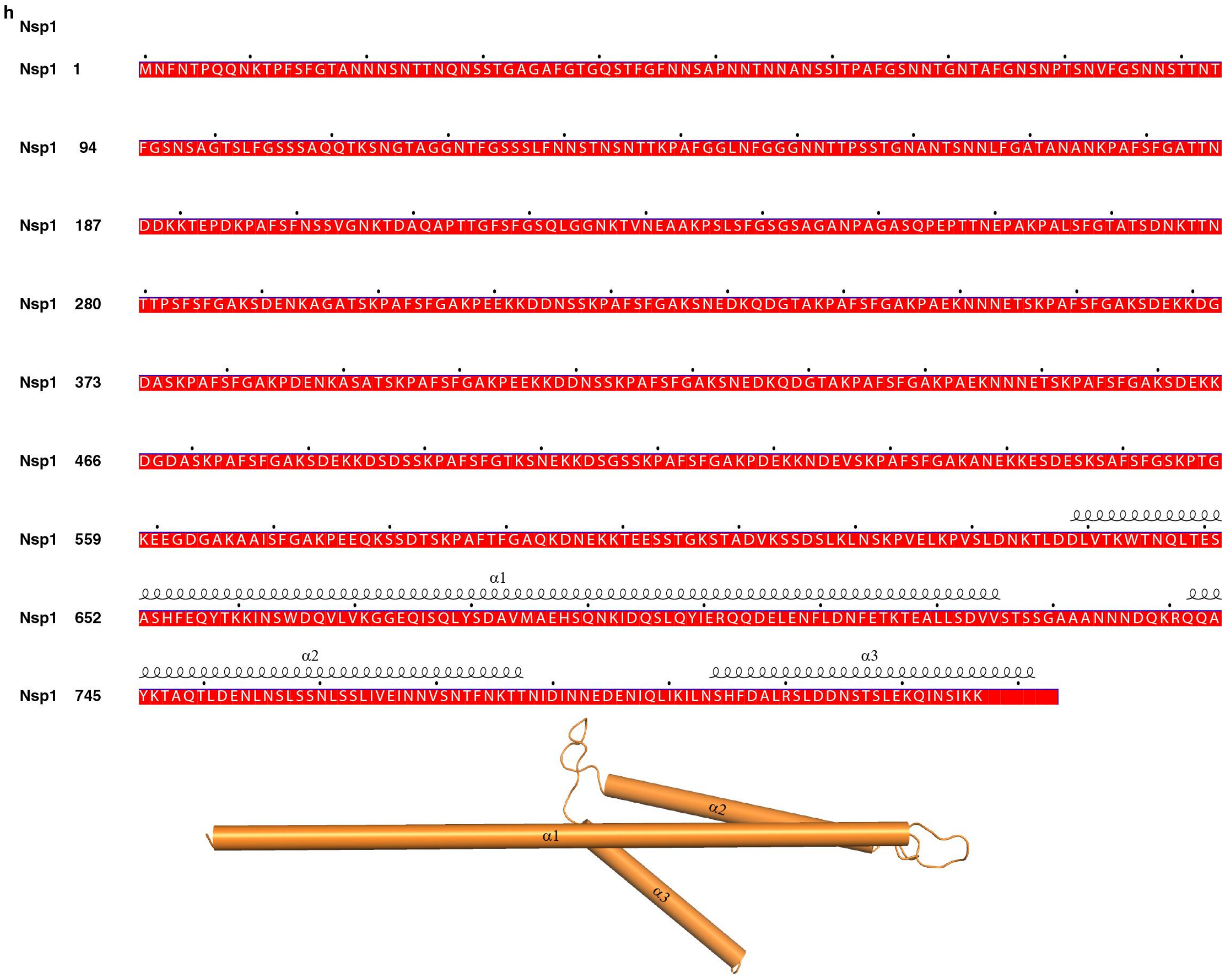


**Supplementary information, Fig. S13. Secondary structural information of IR subunits.** (**a-h**) Secondary structural information of Nup188 (a), Nup192 (b), Nup157 (c), Nup170 (d), Nic96 (e), Nup57 (f), Nup49 (g) and Nsp1 (h).

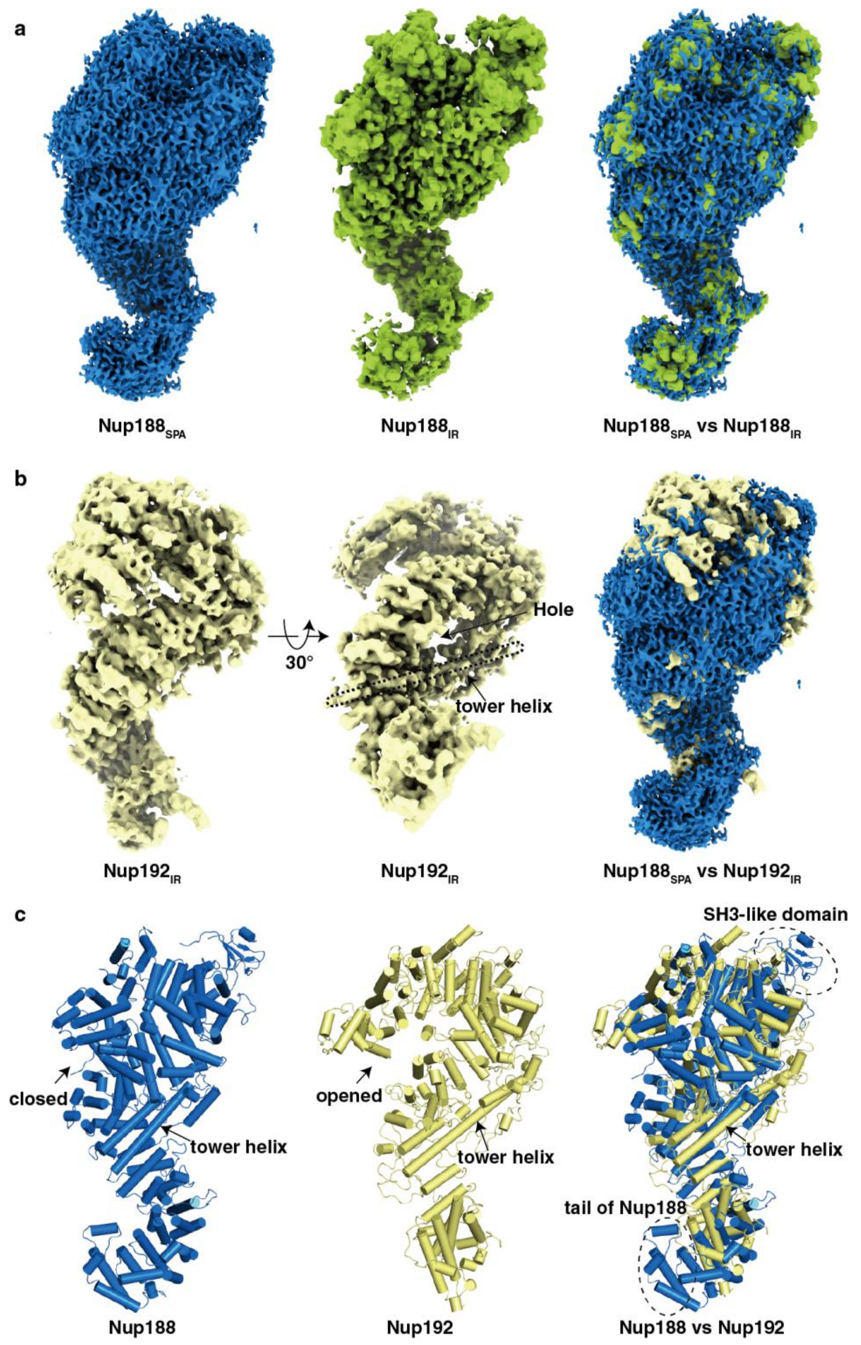


**Supplementary information, Fig. S14. Structures of homologous Nup188 and Nup192.**
(**a**) Comparison of Nup188 maps from single particle analysis (Nup188$_{SPA}$) and IR monomer map (Nup188$_{IR}$). The superposition of Nup188$_{SPA}$ and Nup188$_{IR}$ is shown in the right panel. (**b**) Comparison of density maps of homologous Nup188 and Nup192. Nup192$_{IR}$ represents Nup192 map from IR monomer map. The superposition of Nup188$_{SPA}$ and Nup192$_{IR}$ is shown in the right panel. (**c**) Comparison of atomic models of Nup188 and Nup192. The Nup188 model is built based on the map of Nup188$_{SPA}$ and Nup192 model is generated by homologous modeling based on crystal structure of Nup192 from *Chaetomium thermophilum* (PDB: 5HB4).

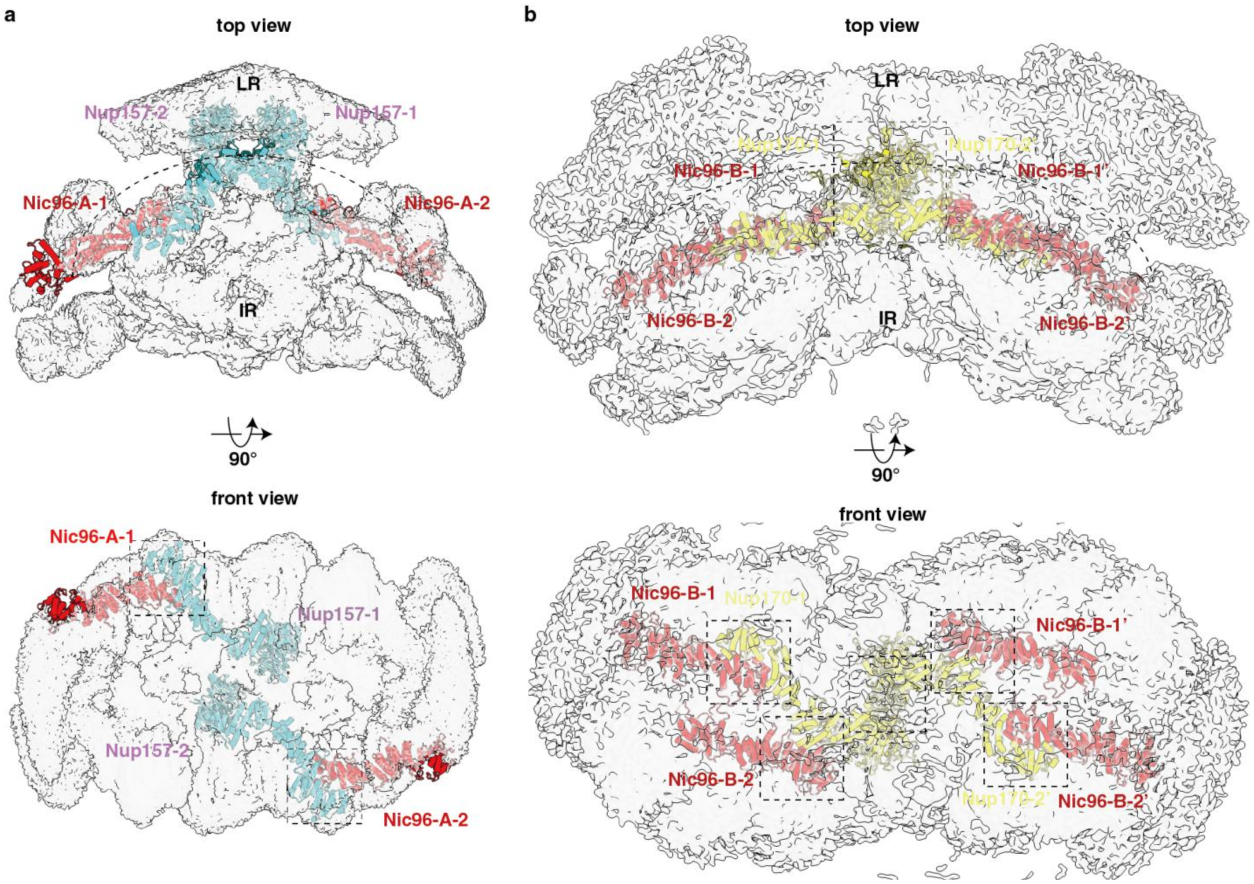


**Supplementary information, Fig. S15. Nup157 and Nup170 mediate conjunction between IR and LR.**

Nup157 (**a**) and Nup170 (**b**) mediate the conjunction between IR and LR by protruding their N-terminals into LR and C-terminal interactions with Nic96 from IR. Dotted curves indicate boundary of LR and IR. Two views are shown and maps of IR monomer and dimer are shown at a transparency of 70%. Abbreviations: LR, luminal ring; IR, inner ring.

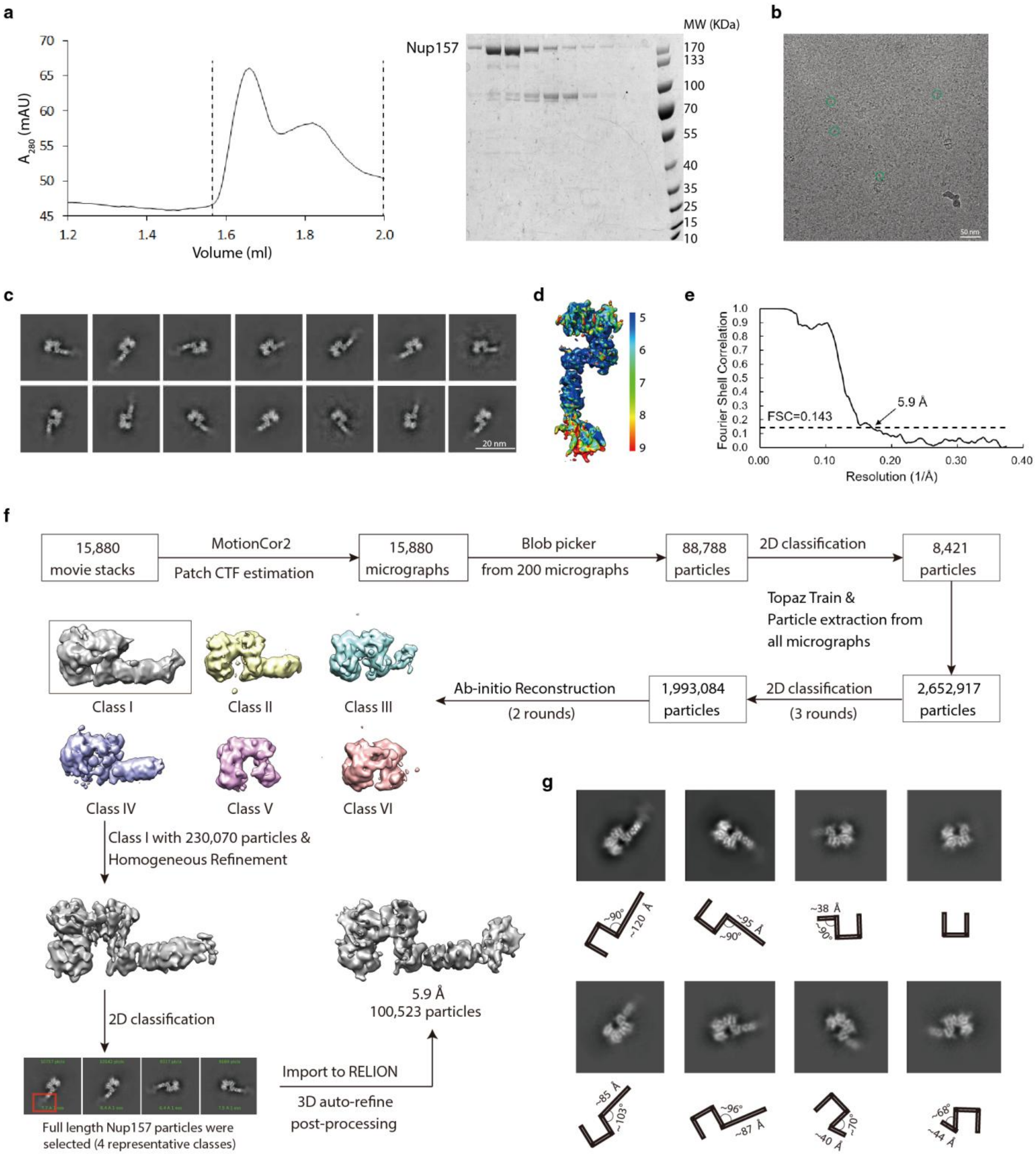


**Supplementary information, Fig. S16. Cryo-EM data analysis of Nup157.**
(**a**) Purification of Nup157. SEC profile of Nup157 (left panel) and the SDS-PAGE gel of the fractions corresponding to the region between dashed lines on SEC curve (right panel). (**b**) A representative raw cryo-EM image for Nup157 with typical particles marked by green circles. (**c**) Typical good reference-free 2D class averages of Nup157. (**d**) Local resolution of cryo-EM map for Nup157. (**e**) Gold standard FSC curves for the cryo-EM maps of whole Nup157. (**f**) The flowchart for EM data processing and the local resolution maps. Details can be found in "Materials and Methods". (**g**) 2D class average images showing the flexible C-terminal of Nup157.

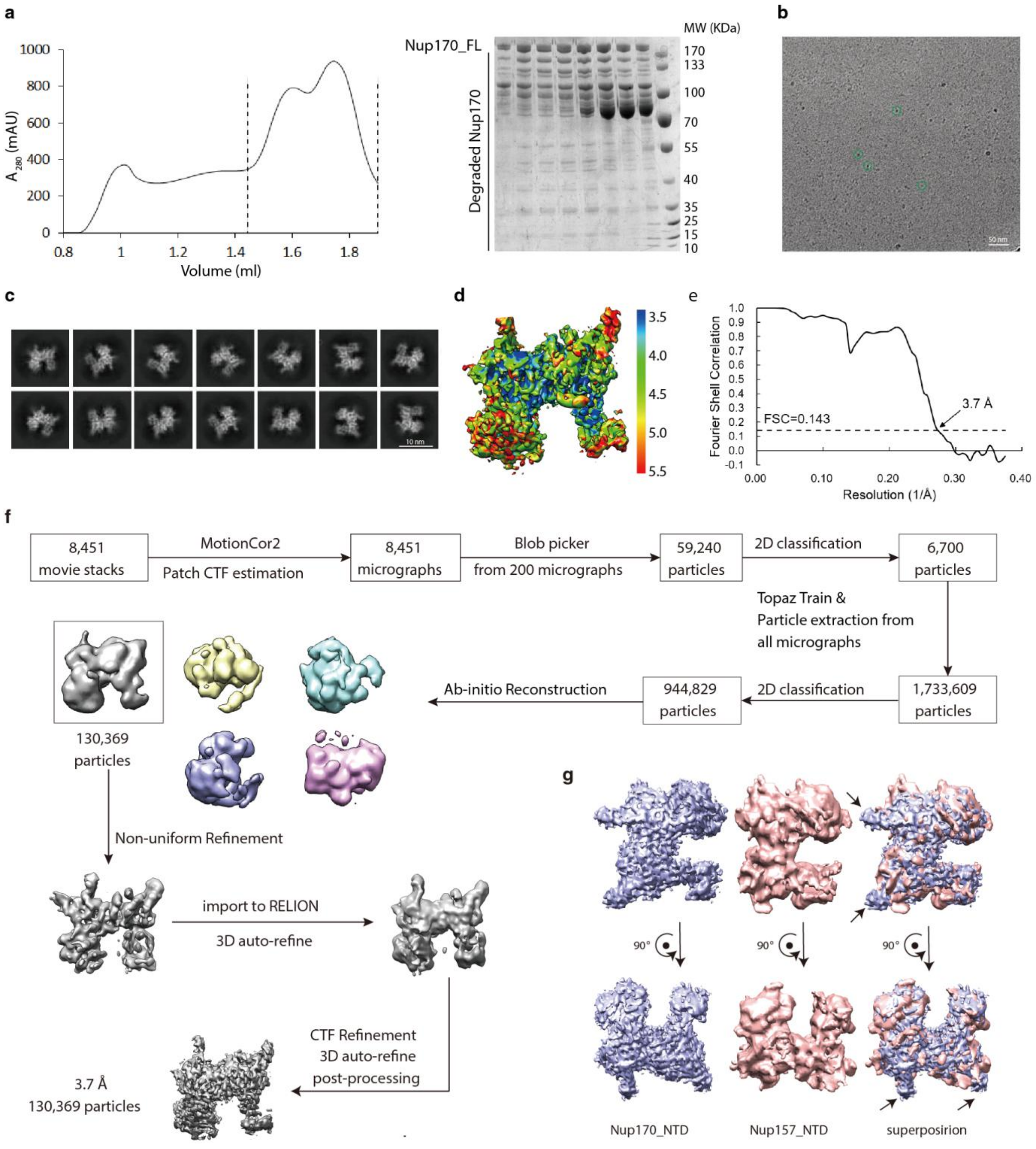


**Supplementary information, Fig. S17. Cryo-EM data analysis of Nup170.**
(**a**) Purification of Nup170. SEC profile of Nup170 (left panel) and the SDS-PAGE gel of the fractions corresponding to the region between dashed lines on SEC curve (right panel). (**b**) A representative raw cryo-EM image for Nup170 with typical particles marked by green circles. (**c**) Typical good reference-free 2D class averages of Nup170. (**d**) Local resolution of cryo-EM map for Nup170. (**e**) Gold standard FSC curves for the cryo-EM maps of whole Nup170. (**f**) The flowchart for EM data processing and the local resolution maps. Details can be found in "Materials and Methods". (**g**) Comparison of the density map of the N-terminal domain (NTD) of Nup157 and Nup170. The superposition is shown in the right panel.

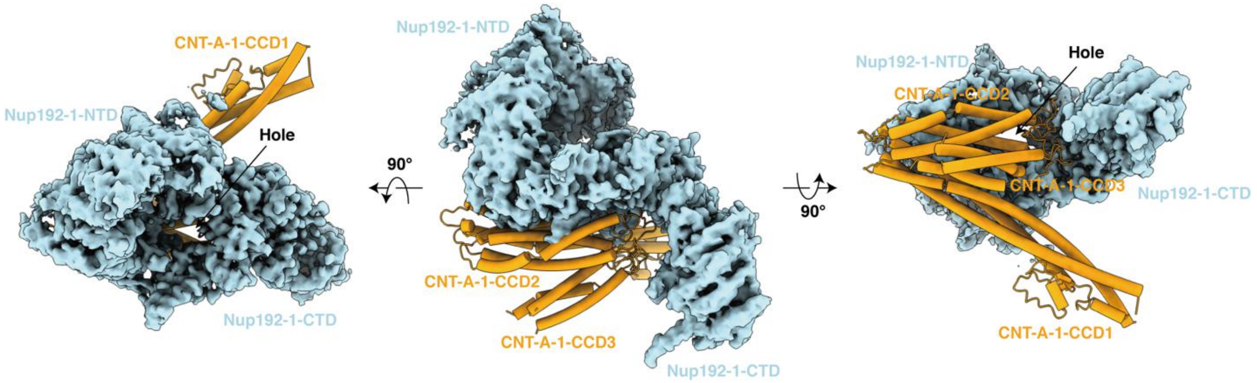


**Supplementary information, Fig. S18. Interaction between Nup192 and CNT.**
The loops linking CCD2 and CCD3 of CNT-A-1 protrude into the C-terminal of Nup192-1, which blocks the hole located between N-terminal domain (NTD) and C-terminal domain (CTD) of Nup192.

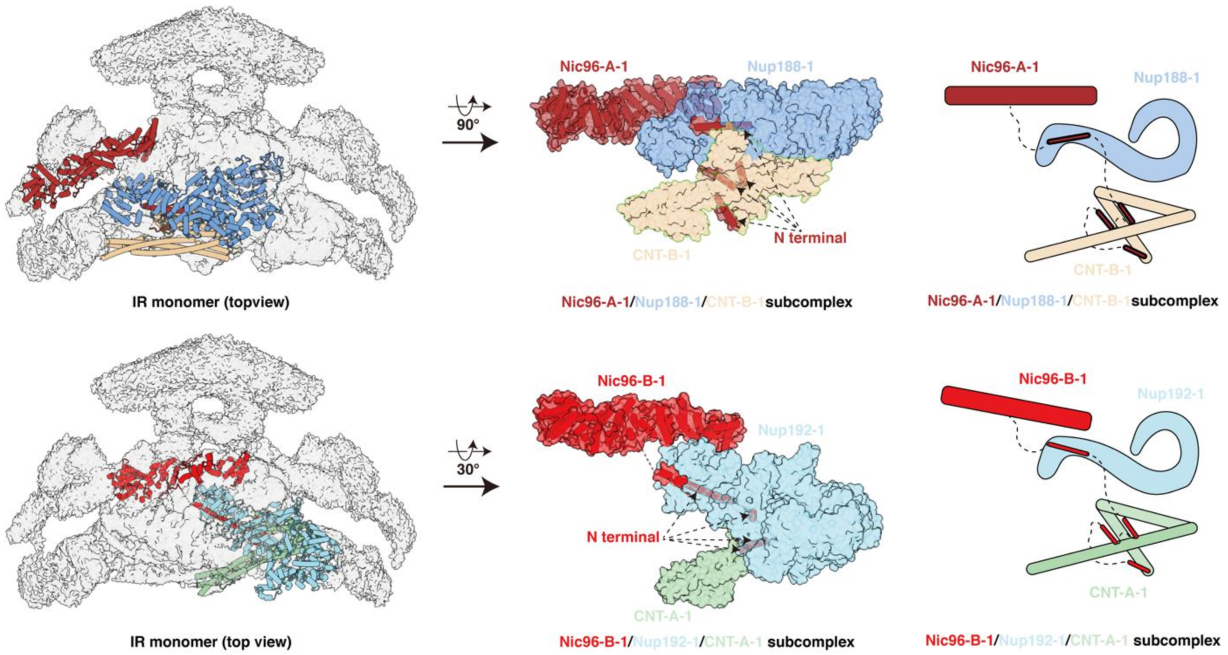


**Supplementary information, Fig. S19. Location of N-terminal α-helices of Nic96 and the density for Nup53/59.**

N-terminal of Nic96 strings three layers of IR monomer through the interaction with Nup188, Nup192 and CNT complex by several α-helices.

Table S1. Plasmids and yeast strains used in this study.

| Plasmids | Expressed strains | Reference |
|---|---|---|
| pESC-His-Nup192-Flag | yeast | This study |
| pESC-Ura-Nup188-Flag | yeast | This study |
| pESC-Leu-Nup157-Flag | yeast | This study |
| pESC-Leu-Nup170-Flag | yeast | This study |
| pGEX-6t-Nup157 | BL21(DE3) | This study |
| pET28at-SUMO-Nup170 | BL21(DE3) | This study |
| pET28at-SUMO-Nup192 | BL21(DE3) | This study |
| pCDNA-Nup157-3xFlag | 293F | This study |
| pCDNA- Nup170-3xFlag | 293F | This study |
| pCDNA- Nup192-3xFlag | 293F | This study |

| Yeast Strains | Genotype | Reference |
|---|---|---|
| W303α | MATα ura3-52 leu2-3,112 his3-11,15 trp1 | Wild type |
| W303a | MATa ura3-52 leu2-3,112 his3-11,15 trp1 | Wild type |
| W303α-Mlp1-PrA | MATα ura3-52 leu2-3,112 his3-11,15 trp1 Mlp1-TEV-ProteinA::LEU2 | This study |
| W303α-84-3FH/Mlp1-PrA | MATα ura3-52 leu2-3,112 his3-11,15 trp1 Nup84-3x Flag-10x His :: URA3 Mlp1-TEV-ProteinA::LEU2 | This study |
| W303a-Mlp1-PrA | MATa ura3-52 leu2-3,112 his3-11,15 trp1 Mlp1-TEV-ProteinA::LEU2 | This study |
| W303a-84-3FH/Mlp1-PrA | MATa ura3-52 leu2-3,112 his3-11,15 trp1 Nup84-3x Flag-10x His :: URA3 Mlp1-TEV-ProteinA::LEU2 | This study |

**Table S2. Cryo-EM data collection and model statistics of NPC and nucleoporins.**

| | Nup188 | IR monomer | IR protomer | IR dimer | IR | NPC | Nup157 | Nup170 |
|---|---|---|---|---|---|---|---|---|
| **Data collection and processing** | | | | | | | | |
| Magnification | 130,000 | 130,000 | 130,000 | 130,000 | 130,000 | 130,000 | 130,000 | 130,000 |
| Voltage (kV) | 300 | 300 | 300 | 300 | 300 | 300 | 300 | 300 |
| Camera | K3 | K3 | K3 | K3 | K3 | K3 | K3 | K3 |
| Electron exposure ($e^{-}/Å^2$) | 50 | 50 | 50 | 50 | 50 | 50 | 50 | 50 |
| Defocus range (μm) | -1.5 ~ -2.5 | -1.5 ~ -2.5 | -1.5 ~ -2.5 | -1.5 ~ -2.5 | -1.5 ~ -2.5 | -1.5 ~ -2.5 | -1.5 ~ -2.5 | -1.5 ~ -2.5 |
| Pixel size (Å) | 0.668 | 0.668 | 0.668 | 0.668 | 0.668 | 0.668 | 0.668 | 0.668 |
| Micrographs (no.) | 16,527 | 296,820 | 296,820 | 296,820 | 296,820 | 296,820 | 15,880 | 8,451 |
| Initial particle images (no.) | 1,208,727 | 2,238,689 | 1,266,268 | 885,259 | 278,938 | 279,900 | 2,652,917 | 1,733,609 |
| Final particle images (no.) | 607,216 | 633,134 | 1,266,268 | 331,211 | 89,774 | 51,220 | 100,523 | 130,369 |
| Symmetry imposed | C1 | C2 | C1 | C2 | C8 | C8 | C1 | C1 |
| Map resolution (Å) | 2.8 | 3.73 | 3.71 | 7.69 | 9.10 | 12.03 | 5.9 | 3.7 |
| Map sharpening B factor ($Å^2$) | -119.6 | -143.4 | -163.6 | -763.5 | -860 | -1000 | -461 | -254.4 |
| FSC threshold | 0.143 | 0.143 | 0.143 | 0.143 | 0.143 | 0.143 | 0.143 | 0.143 |
| Map resolution range (Å) | 10.0-2.7 | 10.0-3.7 | 10.0-3.6 | 20.0-6.5 | 30.0-8.0 | 35.0-12.0 | 5.0-9.0 | 3.5-5.5 |
| EMDB number | | | | | | | | |
| **Refinement** | | | | | | | | |
| Initial model used | generated in RELION3.1 | | | | | | | |
| Model composition | | | | | | | | |
| Non-hydrogen atoms | 12,743 | 139520 | 69760 | | | | | |
| Protein residues | 1,581 | 17410 | 8705 | | | | | |
| R.m.s. deviations | | | | | | | | |
| Bond lengths (Å) | 0.002 | 0.017 | 0.017 | | | | | |
| Bond angles (°) | 0.489 | 1.674 | 1.674 | | | | | |
| Validation | | | | | | | | |
| MolProbity score | 1.59 | 3.05 | 3.02 | | | | | |
| Clashscore | 5.55 | 35.01 | 32.70 | | | | | |
| Rotamer outliers (%) | 0 | 4.01 | 4.00 | | | | | |
| Cβ outliers (%) | 0 | 0.88 | 0.88 | | | | | |
| Ramachandran plot | | | | | | | | |
| Favored (%) | 95.86 | 89.54 | 89.52 | | | | | |
| Allowed (%) | 4.14 | 9.22 | 9.25 | | | | | |
| Disallowed (%) | 0 | 1.23 | 1.23 | | | | | |
| PDB accession number | | | | | | | | |

**Movie S1.** The motion of flexible C-terminal region of Nup188 indicated by the 2D class average image obtained by EMAN2.

**Movie S2.** The motion of flexible C-terminal region of Nup188 indicated by the 3D density map obtained by RELION.

**Movie S3.** The motion of flexible C-terminal region of Nup157 indicated by the 2D class average image obtained by EMAN2.

**Movie S4.** The molecular architecture of the IR monomer.
